# Depletion of extracellular asparagine impairs self-reactive T cells and ameliorates autoimmunity in a murine model of multiple sclerosis

**DOI:** 10.1101/2025.06.09.658561

**Authors:** Peter Georgiev, Sheila Johnson, Kiran Kurmi, Song-Hua Hu, SeongJun Han, Dillon Patterson, Thao H. Nguyen, Linglin Huang, Dan Liang, Naomi Goldman, Thomas Conway, Hannah Creasey, Jared Rowe, Marcia C. Haigis, Arlene H. Sharpe

## Abstract

Amino acids play critical roles in the activation and function of lymphocytes. Here we show that the non-essential amino acid, asparagine, is essential for optimal activation and proliferation of CD4^+^ T cells. We demonstrate that asparagine depletion at different time points after CD4^+^ T cell activation reduces mitochondrial membrane potential and function. Furthermore, asparagine depletion at specific time points during CD4^+^ T cell differentiation reduces cytokine production in multiple CD4^+^ T cell subsets. In an adoptive transfer model of experimental autoimmune encephalomyelitis (EAE), myelin oligodendrocyte-specific pathogenic T helper 17 cells differentiated under Asn-deficient conditions exhibited reduced encephalitogenic potential and attenuated EAE severity. In a model of EAE induced by active immunization, therapeutic depletion of extracellular Asn significantly reduced disease severity. These results identify asparagine as a key metabolic regulator of the pathogenicity of autoreactive CD4^+^ T cells and suggest that targeting asparagine metabolism may be a novel therapeutic strategy for autoimmunity.

## INTRODUCTION

Naive T cell activation and clonal expansion are bioenergetically demanding processes that require substantial changes in energy and substrate utilization.^1–3^ Following T cell receptor (TCR) stimulation, activated T cells increase their metabolic activity and shift towards the use of anabolic pathways such as aerobic glycolysis, fatty acid synthesis, and mitochondrial biogenesis.^1–5^ This metabolic transition depends on the uptake of extracellular nutrients and is enabled by the rapid upregulation of nutrient transporters.^6,7^ Among these, the uptake of exogenous amino acids is important for sustaining intracellular biosynthetic processes including nucleotide biosynthesis, ATP generation, and nascent protein synthesis.^7–9^

Amino acids are fundamental building blocks for the synthesis of nascent proteins but have additional functions during T cell activation. Amino acids serve as substrates for nucleic acid biosynthesis, support post translational modifications of proteins and are critical for maintaining redox balance.^7–9^ The reliance of activated T cells on the non-essential amino acids glutamine, serine, alanine, and arginine is well established.^10–13^ For example, activated T cells critically depend on uptake of serine, which provides glycerol and carbon units for de novo nucleotide biosynthesis and one-carbon metabolism.^12^ Similarly, uptake of alanine is required for T cell proliferation and the efficient exit from quiescence following TCR stimulation.^13^ Extracellular asparagine (Asn) has recently garnered attention as a nutrient important for CD8^+^ T cell differentiation and function, however its role in helper CD4^+^ T cell responses has not been explored.^14–18^ In fact, a number of extracellular amino acid dependencies have yet to be characterized in the specific context of CD4^+^ T cell function. To address this gap, we assessed the requirement for extracellular non-essential amino acids during CD4^+^ T cell activation and proliferation. Through this analysis, we uncovered a critical role for extracellular Asn in supporting CD4^+^ T cell responses.

In this study, we demonstrate that CD4^+^ T cells depend on extracellular Asn for optimal proliferation, activation, and differentiation into helper cells, despite upregulating the Asn-generating enzyme Asn synthetase (ASNS). The absence of extracellular Asn impairs TCR-induced metabolic reprogramming and results in dysfunctional mitochondria. Our findings reveal that early or delayed Asn depletion during the course of experimental autoimmune encephalomyelitis (EAE) attenuates disease severity. In an adoptive transfer model of EAE, pathogenic T helper 17 cells differentiated in the absence of extracellular Asn accumulate poorly in the central nervous system (CNS) and exhibit defects in protein synthesis and metabolic fitness ex vivo. Collectively, our results suggest that extracellular Asn bioavailability is a key metabolic checkpoint that regulates the functional responses of CD4^+^ T cells.

## RESULTS

### Extracellular asparagine (Asn) is essential for optimal activation and proliferation of CD4^+^ T cells

To investigate the requirement for non-essential amino acids in CD4^+^ T cell activation, we performed experiments under culture conditions where amino acids were individually added or withdrawn from media. For single amino acid addition experiments, naive CD4^+^ T cells were activated on plates coated with anti-CD3 and anti-CD28 antibodies and cultured in DMEM media with glutamine. Non-essential amino acids lacking in standard DMEM formulation, but present in the conventional RPMI formulation, were individually added. After 24 hours of stimulation in these culture conditions, only the addition of Asn was sufficient to fully activate CD4^+^ T cells and promote expression of canonical activation markers (**Figure 1A, D-I**). After 72 hours of activation in these culture conditions, only Asn addition led to significant proliferation, as measured by CTV dye dilution (**Figure 1A-C**). To assess the essentiality of non-essential amino acids for activation of naive CD4^+^ T cells, we activated naive CD4^+^ T cells in RPMI lacking nine non-essential amino acids or RPMI depleted of each non-essential amino acid for 24 hours. Depletion of Asn or glutamine resulted in reduced expression of activation markers CD44, CD25, and PD-1, and phenocopied the effects seen when all nine non-essential amino acids were reduced (**Figure 1J-K, S1A**). However, Asn deprivation resulted in the greatest defect in expression of activation markers compared to other amino acids (**Figure 1J-K, S1A**). To determine the dose dependency of Asn for CD4^+^ T cell activation, we performed a titration experiment in which CD4^+^ T cells were stimulated in Asn-free RPMI or DMEM supplemented with increasing concentrations of asparagine. After 24 hours of stimulation, activation marker expression was measured. The resulting titration curve revealed that the critical Asn concentration required for CD4^+^ T cell activation lies between 3.78 and 37.8 µM (**Figure S1B-C**), consistent with the physiological concentration of asparagine in murine plasma, which is approximately 50 μM.^19–20^ These results show that Asn is essential for CD4^+^ T cell activation in these culture conditions.

**Figure 1.**
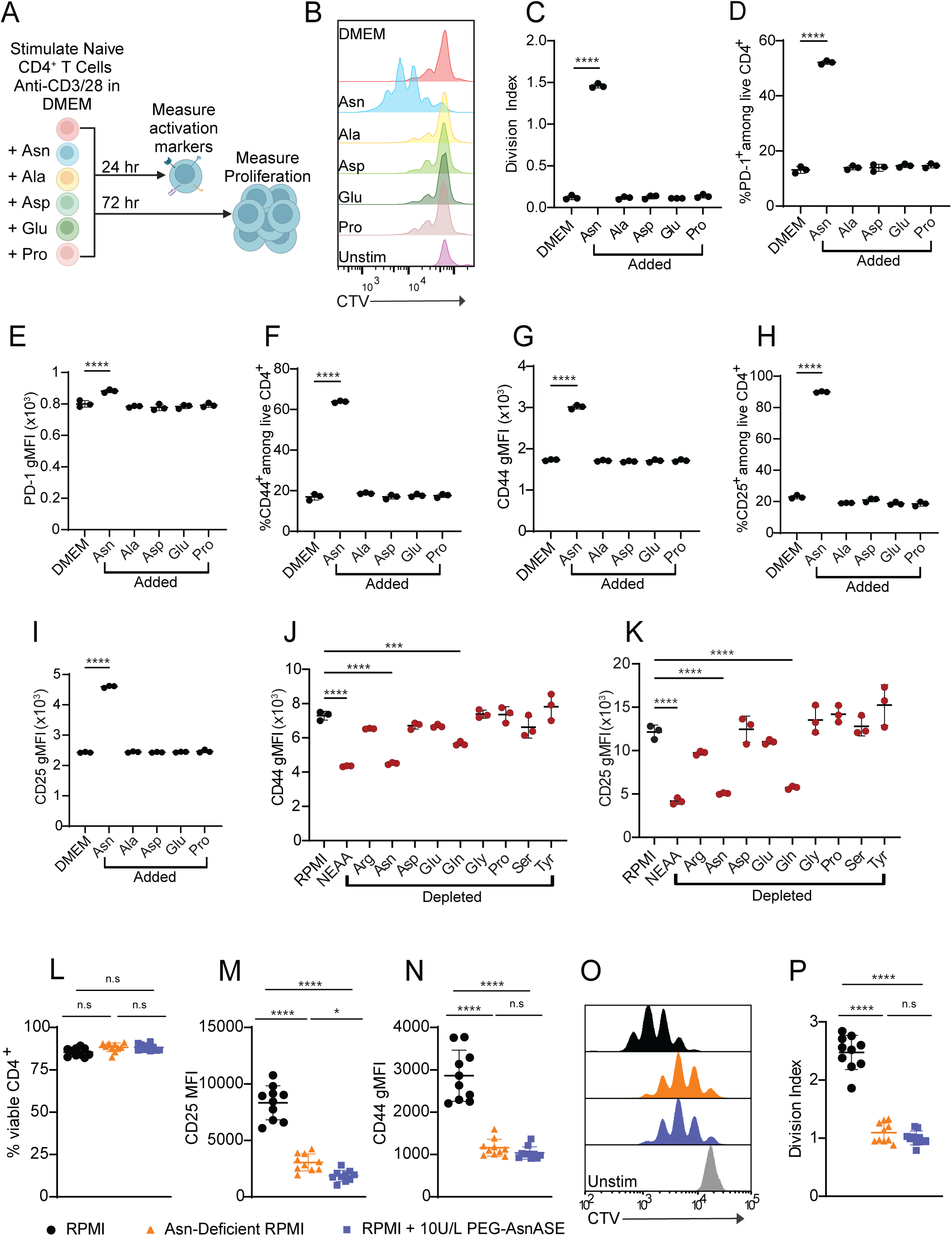
Asparagine is critical for early activation and proliferation of CD4^+^ T cells. (**A**) Schematic of experimental design. Naive CD4^+^ T cells were stimulated for either 24 hours or 72 hours with plate-bound anti-CD3 and anti-CD28 mAbs in DMEM media with glutamine or DMEM media with glutamine supplemented with 0.38 mM asparagine (Asn), 0.38 mM alanine (Ala), 0.15 mM aspartate (Asp), 0.13 mM glutamate (Glu), or 0.17 mM proline (Pro). (**B**) Representative flow cytometry histogram depicting CTV dye dilution in naive CD4^+^ T cells following 3 days of stimulation with plate-bound anti-CD3/CD28 mAbs in DMEM media with glutamine or DMEM media with glutamine supplemented with the indicated amino acids. (**C**) Quantification of division index in B. (**D-I**) Quantification of the proportions of CD4^+^ T cells expressing the cell surface activation markers PD-1 (**D**), CD44 (**F**) CD25 (**H**) as well as expression levels of each respective marker on a per cell basis (**E,G,I**) following 24 hours of stimulation with plate-bound anti-CD3/CD28 mAbs in DMEM media with glutamine or DMEM media with glutamine supplemented with 0.38 mM Asn, 0.38 mM Ala, 0.15 mM Asp, 0.13 mM Glu, or 0.17 mM Pro. (**J-K**) Expression levels of activation markers CD44 (**J**), CD25 (**K**) following 24 hours of stimulation with plate-bound anti-CD3/CD28 mAbs in RPMI lacking the indicated individual amino acids shown in red. Non-essential amino acids (NEAA) include asparagine (Asn), aspartate (Asp), glutamate (Glu), proline (Pro), arginine (Arg), glutamine (Gln), glycine (Gly), serine (Ser), and tyrosine (Tyr). (**L**) Quantification of the proportions of viable CD4^+^ T cells following 24 hours of stimulation with plate-bound anti-CD3/CD28 mAbs in complete RPMI (RPMI), Asn-deficient RPMI or RPMI with 10 IUs/L PEGylated-asparaginase (PEG-AsnASE) added at the start of culture. (**M-N**) Expression level of CD25 (**M**) and CD44 (**N**) on a per cell basis following 24 hours of stimulation with plate-bound anti-CD3/CD28 mAbs under the same conditions as in L. (**O**) Representative flow cytometry histograms showing CTV dye dilution in naive CD4^+^ T cells following 3 days of stimulation with plate-bound anti-CD3/CD28 mAbs in RPMI, Asn-deficient RPMI or RPMI with 10 IUs/L PEG-AsnASE added at the start of culture. Grey histogram represents unstimulated control. (**P**) Quantification of division index in O. Each dot represents cells obtained from an individual animal (L-N, P). Results are shown as mean ± SD (C-N, P) and are representatives of 2 independent experiments (B-K, O) or pooled from 2 independent experiments (L-N, P). Not significant (n.s), *p < 0.05, ***p < 0.001, ****p < 0.0001, one-way ANOVA with Dunnet’s multiple comparison test (C-K) or Tukey’s multiple comparison test (L-N, P).

To further evaluate the extent to which extracellular Asn depletion affects CD4^+^ T cell activation and proliferation, we employed two orthogonal approaches to deplete Asn. We activated naive CD4^+^ T cells in media treated with PEGylated asparaginase (PEG-AsnASE), an enzyme that catabolizes asparagine into aspartate and ammonia, or in custom-formulated RPMI media specifically lacking Asn. Viability did not significantly differ between complete or Asn-deficient media conditions (**Figure 1L**), however, cells cultured in Asn-deficient media displayed reduced expression and frequencies of CD25, CD44, CD69, and PD-1 positive cells after 24 hours in culture (**Figure 1M-N, S1D-I**). These results were mimicked with PEG-AsnASE treated media. In contrast, at longer culture periods after TCR stimulation, cells demonstrated increased apoptosis as indicated by Annexin V+ staining at 48 hours (**Figure S1K-L**) and reduced viability after 72 hours in Asn-deficient media (**Figure S1J**). Proliferation was also impaired by the depletion of Asn after 3 days of TCR stimulation (**Figure 1O-P**). In sum, CD4^+^ T cells exhibit a requirement for extracellular Asn at early stages of activation.

Asparagine is described as a non-essential amino acid and thus can be produced endogenously. Considering the essentiality of extracellular Asn during the early stages of CD4^+^ T cell activation, we next explored whether CD4^+^ T cells lack the intracellular machinery needed for endogenous production of Asn (**Figure 2A**). To evaluate the levels of Asn metabolism proteins present in CD4^+^ T cells, we used a publicly available bulk RNA-seq dataset consisting of naive CD4^+^ T cells differentiated in vitro under T helper 1 (TH1), non-pathogenic T helper 17 (npTH17), and pathogenic T helper 17 (pTH17) polarizing conditions for 1, 6, 12, 20 or 48 hours (**Figure 2B-C**).^21^ These data suggest that the Asn synthesizing enzyme, asparagine synthase (*Asns*), increases in expression upon activation under all polarizing conditions, whereas the asparagine catabolizing enzyme, *Asrgl1*, expression demonstrates little change upon activation. We validated these findings in CD4^+^ T cells activated under non polarizing conditions (**Figure 2D-E**) using quantitative PCR (qPCR). To confirm that the observed transcriptional changes correspond to protein expression levels, we assessed ASNS protein expression by western blotting (**Figure 2F**). The changes in ASNS protein levels reflected the transcriptional changes of *Asns*. Notably, when naive CD4^+^ T cells were activated in Asn-limited media, there was greater production of ASNS compared to cells cultured in Asn sufficient media (**Figure 2G**). Together, these results show that activated CD4^+^ T cells possess the enzymatic machinery to generate Asn in both Asn-depleted and Asn-replete conditions.

**Figure 2.**
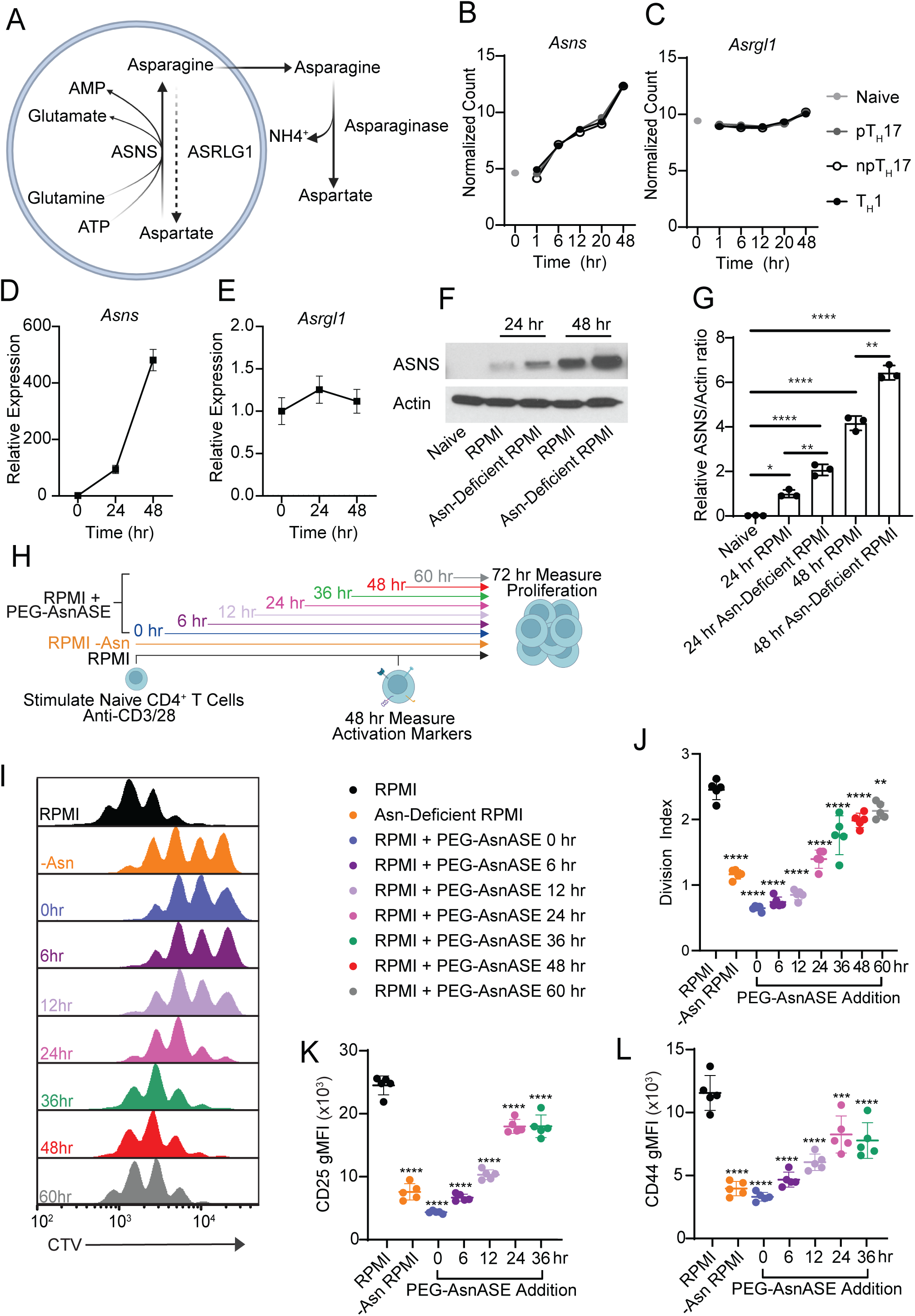
Extracellular asparagine is needed for sustained CD4^+^ T cell activation and proliferation at later stages following initial activation. (**A**) Schematic of Asn metabolism in mammalian cells. (**B-C**) Bulk RNA-seq analysis showing expression of *Asns* (**B**), and *Asrgl1* (**C**) in naive CD4^+^CD62L^+^CD44^-^ T cells at baseline (gray dot) or following stimulation with anti-CD3/CD28 mAbs under T helper 1 (T_H_1), pathogenic T helper 17 (pT_H_17) and non-pathogenic T helper 17 (npT_H_17) conditions for 1, 6, 12, 20 and 48 hours. Results are shown as average (n=3 for each condition, per time point). (**D-E)** qPCR analysis showing the expression kinetics of *Asns* (**D**) and *Asrgl1* (**E**) over time in naive CD4^+^ T cells at baseline or stimulated with anti-CD3/CD28 mAbs for 24 and 48 hours. (**F**) Western blot analysis of ASNS protein expression in CD4^+^ T cells activated with anti-CD3/CD28 mAbs in either RPMI or Asn-deficient RPMI at 24 or 48 hours after activation. Naive CD4^+^ T cells are shown as controls. (**G**) Densitometry analysis of F showing the relative Asns to actin ratio. (**H**) Schematic of experimental design. Purified naive CD4^+^ T cells were activated in vitro with plate-bound anti-CD3/CD28 mAbs in complete RPMI media (RPMI), Asn-deficient RPMI, or RPMI with 10 IUs/L PEGylated-asparaginase (PEG-AsnASE) added at 0, 6, 12, 24, 36, 48 and 60 hours. (**I**) Representative flow cytometry histogram showing cell trace violet (CTV) dye dilution in naive CD4^+^ T cells following 3 days of stimulation. (**J**) Quantification of division index in I. (**K-L**) Quantification of the gMFI of CD4^+^ T cells expressing CD25 (**K**) and CD44 (**L**) following 2 days of stimulation with plate-bound anti-CD3/CD28 mAbs in complete RPMI, Asn-deficient RPMI or RPMI with 10 IUs/L PEG-AsnASE added at 0, 6, 12, 24, 36 hours. Each dot represents cells from an individual animal (J-L). Results are shown as average (n=3 per condition, per time point) (B-C) or mean ± SD (D-E, G, J-L) and are representative of 2 independent experiments (D-G, I-L). non-significant (n.s.), *p < 0.05, **p < 0.01, ***p < 0.001, ****p < 0.0001, one-way ANOVA with Dunnet’s multiple comparison test (J-L) or Tukey’s multiple comparison test (G).

Given that upregulation of ASNS allows CD8^+^ T cells to function in the absence of extracellular Asn^14,17^ and our findings showing that ASNS is upregulated in CD4^+^ T cells upon TCR activation in the presence and absence of Asn (**Figure 2F-G**), we next investigated whether endogenous Asn could compensate for the lack of extracellular Asn and promote CD4^+^ T cell function at 24 and 48 hours after TCR stimulation. To interrogate when extracellular Asn is needed for CD4^+^ T cell activation and proliferation, we depleted extracellular Asn using PEG-AsnASE at different time points following activation, including 6, 12, 24, 36, 48 and 60 hours post TCR activation (**Figure 2H**). Depletion of extracellular Asn significantly inhibited CD4^+^ T cell proliferation, even when PEG-AsnASE was added to cultures at 48 hours post TCR activation (**Figure 2I-J**). The anti-proliferative effects of PEG-AsnASE treatment were statistically significant at time points corresponding to increased ASNS expression, including 24 and 36 hours after TCR activation (**Figure 2D and 2F-G**). In addition, depletion of extracellular Asn at various time points after TCR activation impaired the activation of naive CD4^+^ T cells, reducing the proportions of CD4^+^ T cells expressing activation-associated markers (**Figure S2A-C**). Expression levels of these markers were also reduced in the absence of Asn and never fully recovered to levels observed in control conditions (**Figure 2K-L**). These results suggest that extracellular Asn availability is not only limiting early during TCR activation, but also at later stages, despite increased ASNS levels.

### Extracellular Asn is a building block for protein synthesis following T cell activation

Because amino acids act as fundamental building blocks for the synthesis of nascent proteins,^8^ we reasoned that deficits in the activation and proliferation of CD4^+^ T cells upon Asn deprivation might result from decreased protein synthesis.^8^ To measure levels of protein synthesis in anti-CD3/anti-CD28 activated CD4^+^ T cells, we used fluorescently labeled O-propargyl-puromycin (OPP), an alkyne analog of puromycin which is directly incorporated into the C-terminus of translating polypeptide chains (**Figure 3A**). Naive CD4^+^ T cells stimulated in Asn-deficient RPMI for 24 hours exhibited a significant deficit in nascent protein synthesis, similar to levels observed in unstimulated controls and control cells treated with the protein synthesis inhibitor cycloheximide (**Figure 3B-C**). Adding Asn back to Asn-deficient RPMI largely restored protein synthesis (**Figure 3B-C**). These results demonstrate a requirement for Asn in tRNA charging to support nascent polypeptide synthesis during the activation of naive CD4^+^ T cells.

**Figure 3.**
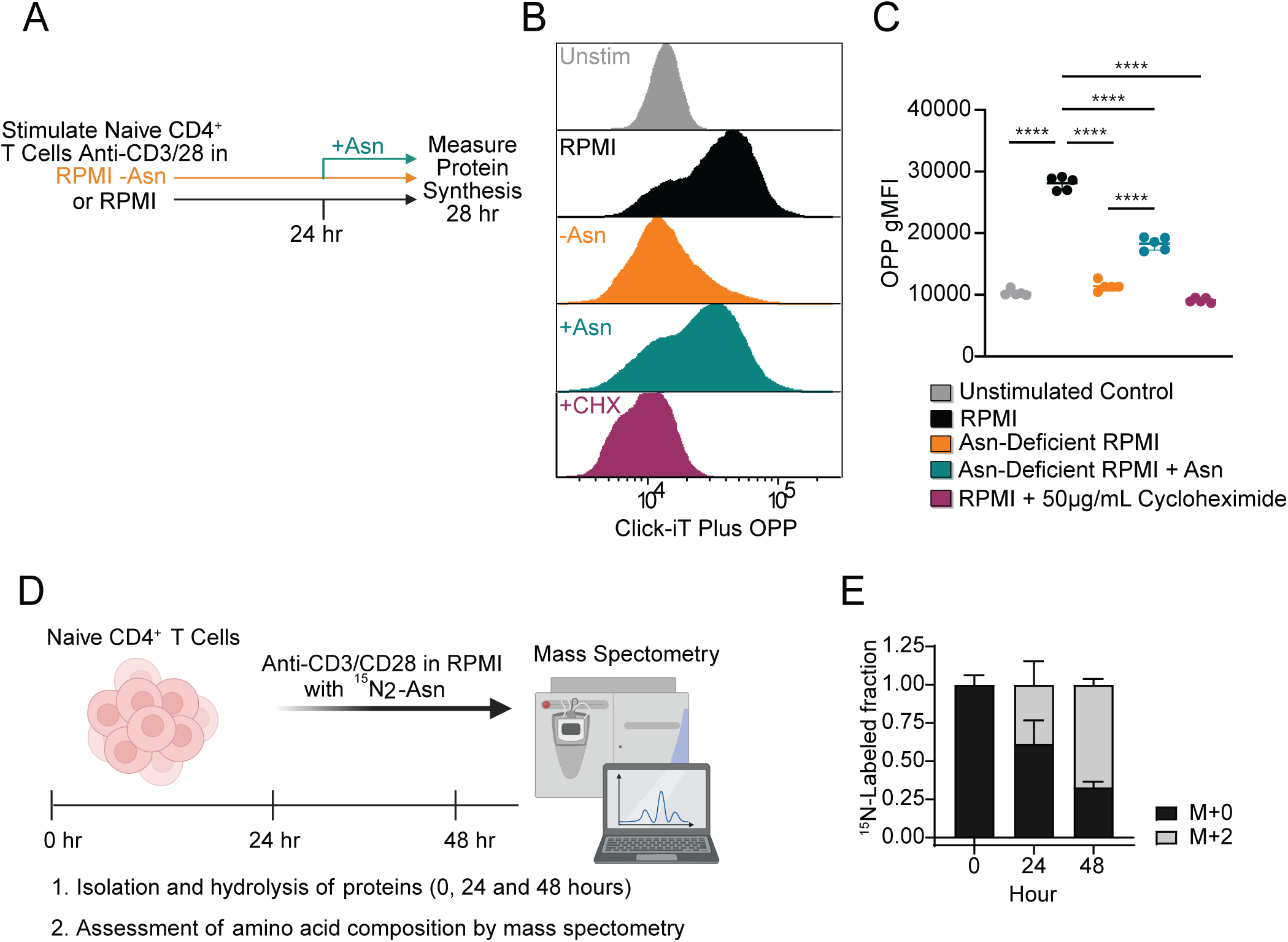
Asparagine is taken up by CD4^+^ T cells and incorporated into their proteome. (**A**) Schematic of experimental design. Purified naive CD4^+^ T cells were activated in vitro with plate-bound anti-CD3/CD28 mAbs in complete RPMI media (RPMI) or Asn-deficient RPMI for 24 hours. At 24 hours Asn was added to cells stimulated in Asn-deficient media at a final concentration of 0.38 mM for 4 hours. At 28 hours protein synthesis was measured using the O-propargyl-puromycin (OPP) probe. As a positive control, T cells cultured in RPMI were treated with 50 μg/mL of the protein synthesis inhibitor cycloheximide (CHX). (**B**) Representative flow cytometry histogram showing mean fluorescent intensity (MFI) of the nascent protein synthesis reporter Click-iT OPP. Naive CD4^+^ T cells are shown as controls. (**C**) Quantification of Click-iT OPP gMFI. (**D**) Experimental design for measuring the incorporation of heavy labeled ^15^N_2_-Asn into proteins following naive CD4^+^ T cell activation. Naive CD4^+^ T cells were stimulated with plate-bound anti-CD3/CD28 mAbs in Asn-deficient media reconstituted with 0.38 mM ^15^N_2_-Asn for 24 or 48 hours. (**E**) Quantification of the ^15^N_2_-Asn labeled fraction in the T cell proteome. Results are shown as mean ± SD (C and E) and are representative of at least two independent experiments (B-C, E). ****p < 0.0001, one-way ANOVA with Tukey’s multiple comparison test (C).

Next, we conducted a time course activation study using ^15^N_2_-Asn to assess the contribution of extracellular Asn to total CD4^+^ T cell protein content (**Figure 3D**). LC/MS analysis of hydrolyzed total protein fractions from naive CD4^+^ T cells at baseline, 24 hours, and 48 hours post TCR stimulation revealed that nearly 50% of Asn from proteins was ^15^N_2_ labeled after 24 hours and roughly 75% of Asn was labeled by 48 hours (**Figure 3E**). Collectively, our studies demonstrate that Asn is taken up by CD4^+^ T cells and plays a requisite role in protein synthesis.

Since we observed increased incorporation of extracellular asparagine into the CD4^+^ T cell proteome, we aimed to identify amino acid transporters that could mediate this process. Plasma membrane transporters, Slc38a9 and Slc1a5 have been linked to Asn uptake in activated CD8^+^ T cells^15^, and Slc6a14 has been implicated in Asn uptake in macrophages^22^. Thus, we sought to determine if these transporters are expressed throughout the activation and differentiation of CD4^+^ T cells. Reanalysis of the abovementioned RNA-seq dataset, which includes transcript levels of naive CD4^+^ T cells differentiated under various polarizing conditions, revealed that Slc1a5 increases expression upon activation in all subsets. Similarly, Slc38a2 expression increases 1 hour after activation but subsequently returns to basal levels comparable to those in the naive state across all polarizing conditions. Slc6a14 exhibited lower basal expression in naive cells relative to the other transporters examined, and its expression decreased progressively over the course of differentiation in all CD4^+^ T cell subsets (**Figures S3A-C**). Together, these results indicate that CD4^+^ T cells express Asn transporters capable of mediating Asn uptake and its incorporation into the proteome.

### Asparagine depletion impairs metabolic reprogramming associated with CD4^+^ T cell activation

Following TCR activation, T cells rapidly upregulate aerobic glycolysis and mitochondrial biogenesis to support the bioenergetic demands required for clonal expansion.^1^ Because nutrient uptake and metabolic reprogramming are highly coordinated processes,^2,6^ we next investigated whether extracellular Asn availability affects CD4^+^ T cell bioenergetics. To probe the functional state of mitochondria under conditions of Asn deprivation, we utilized the metabolic dyes Tetramethylrhodamine methyl ester (TMRM) to label active mitochondria with intact membranes and mitotracker green (MTG) to assess mitochondrial mass. (**Figure 4A**). TMRM and MTG fluorescence were decreased following Asn deprivation or treatment with PEG-AsnASE (**Figure S4A-B**). In addition, a population of depolarized mitochondria displaying high MTG staining (referred to as TMRM/MTG low) emerged (**Figure 4B**). TMRM/MTG low cells have previously been shown to mark dysfunctional mitochondria in exhausted T cells.^23^ The proportion of TMRM/MTG low cells increased in the absence of extracellular Asn (**Figure 4B-C**), suggesting that depletion of extracellular Asn supports the accumulation of depolarized mitochondria with impaired fitness.

**Figure 4.**
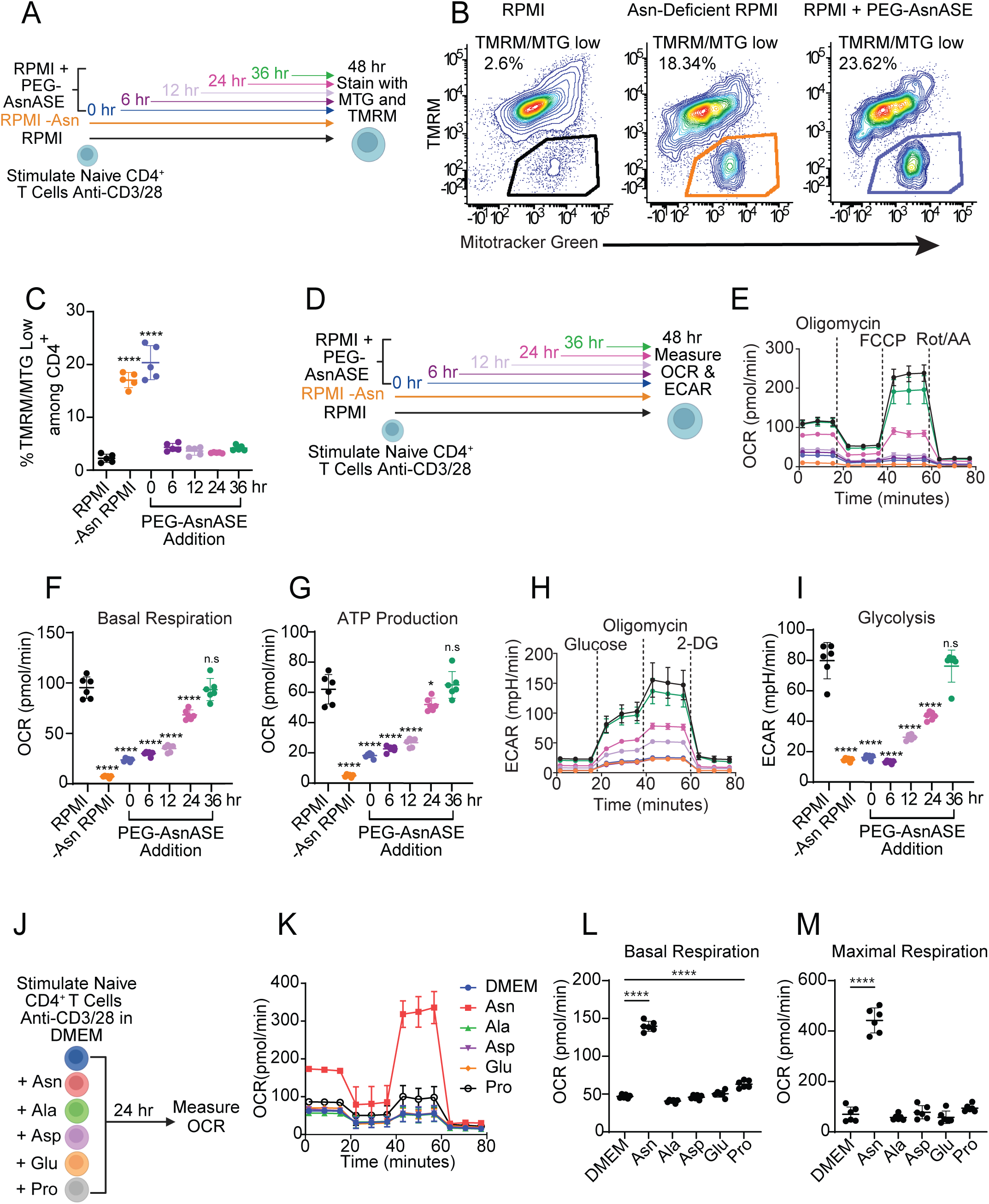
Extracellular asparagine depletion impairs CD4^+^ T cell bioenergetics upon activation. (**A**) Schematic of experimental design. Naive CD4^+^ T cells were stimulated for 48 hours with plate-bound anti-CD3/CD28 mAbs in either complete RPMI media (RPMI), Asn-deficient RPMI or RPMI treated with 10 IUs/L PEGylated-asparaginase (PEG-AsnASE) added at 0, 6, 12, 24, or 36 hours and stained with Mitotracker green (MTG) and Tetramethylrhodamine methyl ester (TMRM). (**B**) Representative flow cytometry contour plots depicting TMRM and MTG in CD4^+^ T cells. (**C**) Quantification of proportion of TMRM/MTG low cells. (**D**) Schematic of experimental design. Oxygen consumption rate (OCR) and extracellular acidification rate (ECAR) were measured in naive CD4^+^ T cells stimulated for 48 hours with plate-bound anti-CD3/CD28 mAbs in either complete RPMI media (RPMI), Asn-deficient RPMI or RPMI with 10 IUs/L PEG-AsnASE added at 0, 6, 12, 24, or 36 hours. (**E**) OCR under mitochondrial stress test conditions. (**F**) Quantification of basal respiration and (**G**) ATP production. (**H**) ECAR under glycolysis stress test conditions. (**I**) Quantification of glycolysis. (**J**) Schematic of experimental design. OCR was measured in naive CD4^+^ T cells stimulated for 24 hours with plate-bound anti-CD3/CD28 mAbs in DMEM with glutamine or DMEM with glutamine supplemented with 0.38 mM of either asparagine (Asn), alanine (Ala), aspartate (Asp), glutamate (Glu), or proline (Pro). (**K**) OCR under mitochondrial stress test conditions. (**L**) Quantification of basal respiration and (**M**) maximal respiration. Each dot in panels C, F-G, and I represent cells obtained from an individual animal. Results are shown as mean ± SD and are representative of at least 2 independent experiments. non-significant (n.s.), *p < 0.05, ****p < 0.0001, one-way ANOVA with Dunnet’s multiple comparison test.

Given our results showing that asparagine deprivation results in the accumulation of dysfunctional mitochondria, we next sought to understand how Asn availability affects overall T cell metabolic reprogramming by measuring mitochondrial respiration and glycolytic flux (**Figure 4D**). Oxidative phosphorylation (OXPHOS), as measured by the oxygen consumption rate (OCR), was significantly decreased in the absence of extracellular Asn under mitochondrial stress test conditions (**Figure 4E**). PEG-AsnASE treatment at 6, 12 or 24 hours post TCR activation led to decreased OCR and ATP production, while PEG-AsnASE treatment at 36 hours post activation did not blunt mitochondrial respiration (**Figure 4E-G**). These results suggest that continuous Asn availability during the early stages of TCR activation is necessary to promote mitochondrial respiration. Under glycolysis stress test conditions, depletion of extracellular Asn similarly reduced the extracellular acidification rate (ECAR), resulting in decreased glycolysis (**Figure 4H-I**). These findings indicate that extracellular Asn availability determines the extent of TCR-induced glycolytic flux and mitochondrial respiration during CD4^+^ T cell activation. Complementary studies using DMEM media supplemented with Asn showed a significant increase in OCR under mitochondrial stress test conditions, resulting in heightened basal respiration and maximal respiration (**Figure 4J-M**). In contrast, DMEM media supplemented with alanine, aspartate, glutamate, or proline failed to sufficiently increase OCR under mitochondrial stress test conditions (**Figure 4J-M**). These results suggest that extracellular Asn is crucial for mitochondrial respiration following T cell activation.

### Asparagine depletion reduces CD4^+^ helper T cell lineage-specific cytokine production

Given the key roles of mitochondrial respiration and amino acids in the specification of distinct helper T cell lineages, ^24–28^ we next tested whether extracellular Asn availability influences the differentiation of naive CD4^+^ T cells into distinct subsets. We differentiated naive CD4^+^ T cells under TH1, npTH17, or pTH17 polarizing conditions, depleting Asn at the initiation of culture or at 0, 12, 24, 36 and 48 hours after activation (**Figure 5A**). Asn depletion impaired the differentiation of all helper T cell subsets tested, whether it was depleted at the initiation of each culture or at specific time points following T cell activation by using PEG-AsnASE (**Figure 5B-J**). Transcription factors defining specific CD4^+^ T cell subsets were reduced when Asn was depleted at the initiation of culture (**Figure S5A-E**). Naive CD4^+^ T cells differentiated under TH1 polarizing conditions exhibited reduced intracellular expression of the TH1-defining cytokine IFN-γ and a reduced proportion of IFN-γ-expressing T cells (**Figure 5C-D**). Asn depletion similarly affected npTH17 and pTH17 cells in the early stages of activation, as intracellular IL-17A expression levels were significantly reduced within both subsets at 0, 12, and 24 hour timepoints (**Figure 5F-G, I-J**). Notably, pTH17 intracellular IL-17A production was specifically affected by PEG-AsnASE treatment in the later timepoints of differentiation compared to npTH17 in these culture conditions (**Figure 5F-G, I-J**). This finding may point to differences in nutrient requirements as naive CD4^+^ T cells differentiate into distinct subsets. These results suggest that Asn depletion impairs the production of lineage-defining cytokines in CD4^+^ helper T cell subsets.

**Figure 5.**
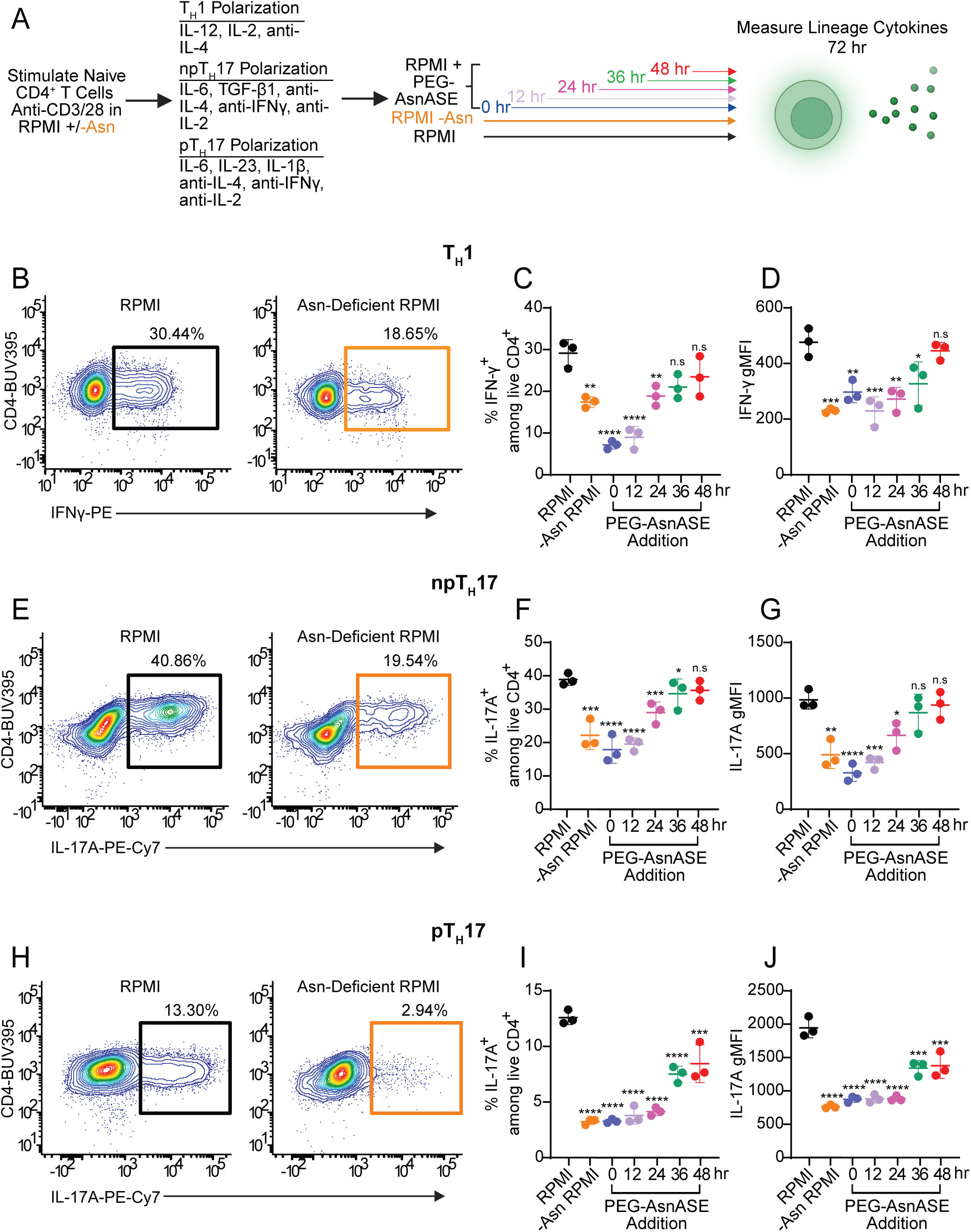
Asparagine deficiency reduces lineage-specific cytokine production in CD4^+^ T helper subsets. (**A**) Schematic of experimental design. Purified naive CD4^+^ T cells were activated in vitro with plate-bound anti-CD3/CD28 mAbs under T helper 1 (T_H_1), pathogenic T helper 17 (pT_H_17), and non-pathogenic T helper 17 (npT_H_17) conditions in either complete RPMI (RPMI) media, Asn-deficient RPMI or RPMI with 10 IUs/L PEGylated-asparaginase (PEG-AsnASE) added at 0, 12, 24, 36 or 48 hours. On day 3 cells were restimulated for 4 hours with phorbol 12-myristate 13-acetate (PMA), ionomycin, brefeldin A and monensin for intracellular staining. (**B**) Representative flow cytometry contour plots depicting intracellular staining of IFN-γ in T_H_1 differentiation conditions in RPMI or RPMI without Asn. (**C**) Quantification of the proportions of IFNγ-producing CD4^+^ T cells under T_H_1 differentiation conditions. (**D**) Quantification of IFN-γ gMFI as shown in C. (**E**) Representative flow cytometry contour plots depicting intracellular staining of IL-17A in npT_H_17 differentiation conditions in RPMI or RPMI without Asn. (**F**) Quantification of the proportions of IL-17A-producing CD4^+^ T cells under npT_H_17 differentiation conditions. (**G**) Quantification of IL-17A gMFI as shown in F. (**H**) Representative flow cytometry contour plots depicting intracellular staining of IL-17A in pT_H_17 differentiation conditions in RPMI or RPMI without Asn. (**I**) Quantification of the proportions of IL-17A-producing CD4^+^ T cells under pT_H_17 differentiation conditions. (**J**) Quantification of IL-17A gMFI as shown in I. Each dot in panels C-D, F-G, and I-J represents cells obtained from an individual animal. Results are shown as mean ± SD and are representative of at least 2 independent experiments. non-significant (n.s.), *p < 0.05 **p < 0.01, ***p < 0.001, ****p < 0.0001, one-way ANOVA with Dunnet’s multiple comparison test.

### Asn depletion ameliorates the severity of experimental autoimmune encephalomyelitis

We hypothesized that by exploiting the dependency of CD4^+^ T cells on extracellular Asn by systemically depleting Asn bioavailability, we could modulate the severity CD4^+^ T cell mediated pathologies, such as experimental autoimmune encephalomyelitis (EAE). To do this, we first prophylactically administered a single dose of PEG-AsnASE to mice one day before inducing EAE through immunization with an emulsion of myelin oligodendrocyte glycoprotein (MOG) peptide, MOG_35–55_, in complete Freund’s adjuvant (CFA) followed by administration of pertussis toxin. Strikingly, while PBS treated controls developed EAE peaking around day 16, PEG-AsnASE treated mice exhibited significantly milder disease with delayed onset (**Figure 6A-C**). To evaluate the therapeutic potential of PEG-AsnASE in a more clinically relevant scenario, we delayed PEG-AsnASE treatment until day 8 after active immunization with MOG_35–55_ (**Figure 6D**). Remarkably, a single dose of PEG-AsnASE was sufficient to attenuate disease severity and delay onset, similar to prophylactic PEG-AsnASE treatment, resulting in a substantially milder disease (**Figure 6D-F**). These results suggest that therapeutic Asn depletion has the potential to be an immunosuppressive strategy to target CD4^+^ T cell-mediated autoimmune pathologies.

**Figure 6.**
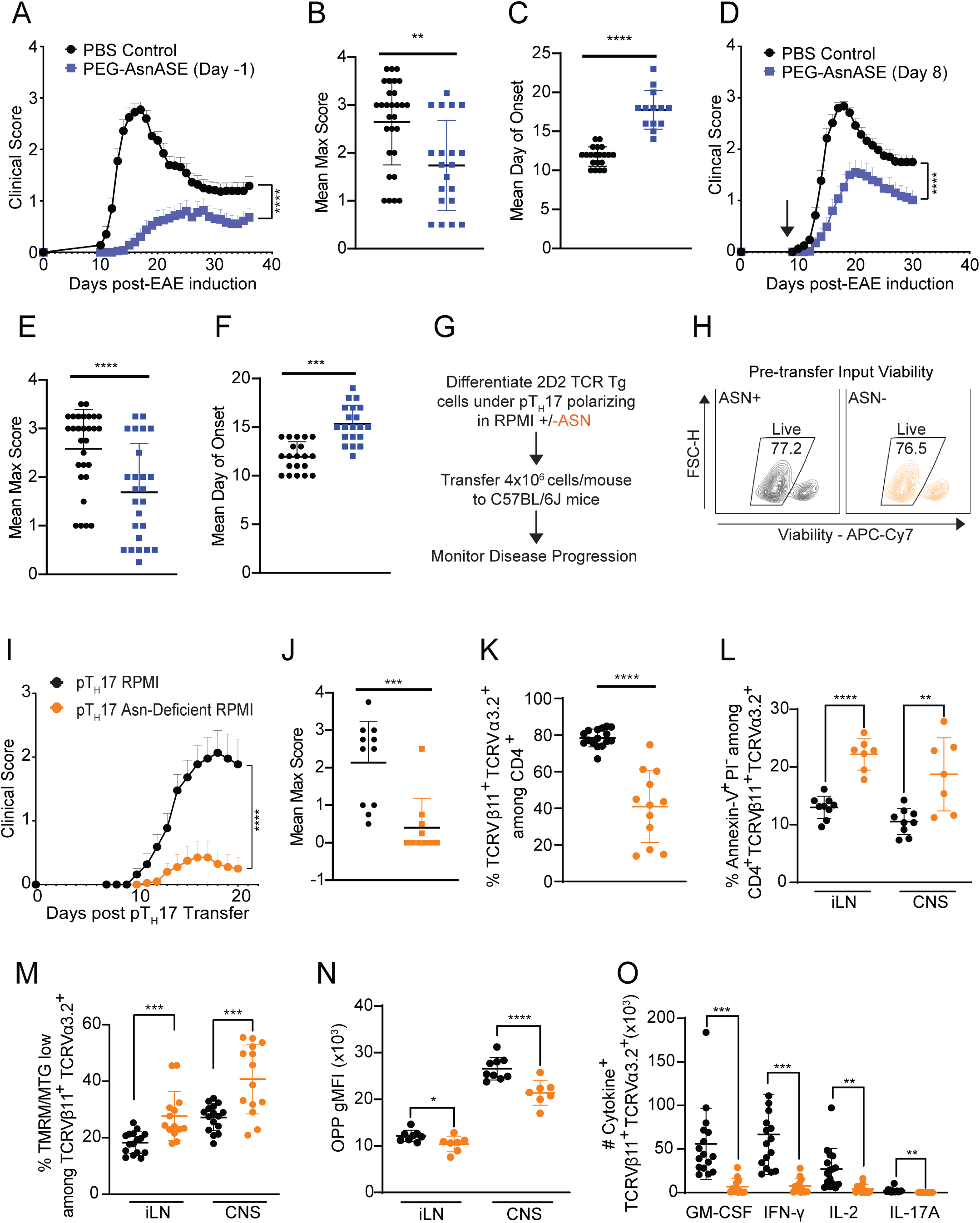
Asparagine deficiency ameliorates experimental autoimmune encephalomyelitis (EAE). (**A**) Mice were treated with a single dose of 25 IUs of PEGylated-asparaginase (PEG-AsnASE) or PBS i.p. one day prior to immunization with MOG_35-55_/CFA and pertussis toxin (PTX) to induce EAE and monitored daily for signs of disease (PBS Control n=20, PEG-AsnASE n=20). (**B**) Quantification of the average maximal EAE scores in PBS vs. PEG-AsnASE treated mice. (**C**) Quantification of the mean day of onset in PBS vs. PEG-AsnASE treated mice. (**D**) EAE was induced by immunization with MOG_35-55_/CFA and pertussis toxin (PTX) and scored daily for disease. Mice were treated with a single dose of 25 IUs of PEG-AsnASE or PBS i.p. on day 8 of active EAE (PBS Control n=20, PEG-AsnASE n=20). (**E**) Quantification of the average maximal EAE scores in PBS vs. PEG-AsnASE treated mice. (**F**) Quantification of the mean day of onset in PBS vs. day 8 PEG-AsnASE treated mice. (**G**) Schematic of experimental design. Pathogenic T helper 17 (pT_H_17) cells were differentiated from naive CD4^+^ FoxP3^-^ T cells from 2D2 TCR transgenic mice in RPMI media with or without Asn and viable 2D2 cells were adoptively transferred (4×10^6^/mouse) into 10-week-old C57BL/6J female recipients to induce EAE. Mice were monitored daily for disease. (**H**) Representative flow plot displaying percentage of viable pT_H_17 polarized 2D2 cells in sufficient and deficient conditions prior to transfer. (**I**) Daily EAE scores (pT_H_17 RPMI n=11, pT_H_17 Asn-deficient RPMI n=10). (**J**) Quantification of the average maximal EAE scores in mice receiving pT_H_17 cells generated in the presence or absence of Asn. (**K**) Quantification of the proportions of CNS-infiltrating Vβ11^+^Vα3.2^+^ 2D2 pT_H_17 cells at peak of EAE (pT_H_17 RPMI n=16, pT_H_17 Asn-deficient RPMI n=12). (**L**) Quantification of the proportions of Vβ11^+^Vα3.2^+^ 2D2 pT_H_17 cells actively undergoing apoptosis (Annexin-V^+^PI^-^) in the CNS and inguinal lymph node at peak of EAE. (**M**) Quantification of the proportions of TMRM/MTG low 2D2 pT_H_17 cells in the CNS and inguinal lymph node at peak of EAE. (**N**) Quantification of OPP gMFI in 2D2 pT_H_17 cells in the CNS and inguinal lymph node at peak of EAE. (**O**) Quantification of the absolute numbers of the indicated cytokines expressed by 2D2 pT_H_17 cells in the CNS at peak of EAE. Results are shown as mean ± SEM (A, D, I) or mean ± SD (B-C, E-F, J-O) and are pooled (A-F, I-K, M, O) or a representative of at least two independent experiments (H, L, N). Each dot represents an individual mouse (B-C, E-F, J-O) *p < 0.05 **p < 0.01, ***p < 0.001, ****p < 0.0001 two-way ANOVA (A, D & I) and Student’s t test (B-C, E-F, and J-O).

### pT_H_17 cells generated in the absence of extracellular Asn are poorly encephalitogenic and exhibit deficits in protein synthesis and mitochondrial fitness in vivo

Since systemic Asn depletion reduces severity of EAE and pTH17 cells are key mediators of EAE, we next investigated how Asn depletion affects the pathogenic potential of pTH17 cells in vivo. We generated pTH17 cells from TCR(Vβ11^+^Vα3.2^+^) transgenic 2D2 mice, which express a TCR specific for myelin oligodendrocyte glycoprotein, using either standard RPMI or Asn-deficient RPMI media and compared their capacity to induce EAE *in vivo*. We adoptively transferred equal numbers of viable 2D2 pTH17 cells into Asn-sufficient WT C67BL/6J hosts and monitored mice for development of EAE (**Figure 6G-H**). 2D2 pTH17 cells generated in Asn-deficient RPMI induced a milder disease compared to 2D2 pTH17 cells generated in RPMI (**Figure 6I-J**). Consistent with these observations, both the proportions and absolute numbers of 2D2 pTH17 T cells from Asn-deficient cultures were significantly reduced in the CNS of recipient mice at the peak of EAE (**Figure 6K, S6A-B**). Furthermore, 2D2 pTH17 cells generated in Asn-deficient RPMI exhibited increased apoptosis, as determined by Annexin V, in the CNS and inguinal lymph nodes (iLN) at peak EAE (**Figure 6L, S6C**). Moreover, 2D2 pTH17 cells generated in Asn-deficient RPMI exhibited reduced MTG and TMRM gMFI in the CNS and iLN (**Figure S6D-E**). There was an enrichment of TMRM/MTG low 2D2 pTH17 cells in the iLN and CNS when differentiated in the absence of Asn (**Figure 6M**), consistent with the deleterious effects of PEG-AsnASE on mitochondrial function in *in vitro* activated CD4^+^ T cells (**Figure 4B-C**).

We next assessed the ex vivo protein synthesis capability of 2D2 pTH17 isolated from the iLN and CNS, using the OPP probe. Both iLN and CNS-infiltrating 2D2 pTH17 differentiated in the absence of Asn exhibited a significant decrease in OPP gMFI at peak of EAE, indicating reduced protein synthesis (**Figure 6N**). The absolute numbers of pathogenic cytokine-producing 2D2 pTH17 T cells generated in Asn-deficient cultures also were reduced in the CNS at peak EAE (**Figure 6O**), further showing the negative impact of Asn deficiency on pTH17 protein synthesis capability. Taken together, these results suggest that the deprivation of extracellular Asn during pTH17 differentiation leads to deficits that reduce the pathogenic potential of autoreactive T cells.

## DISCUSSION

Over the past two decades there have been significant advances in our understanding of the metabolic processes involved in T cell function and lineage commitment. This has increased interest in the potential of modifying T cell responses using targeted metabolic perturbations.^3,29–30^ To meet the metabolic demands of activation and clonal expansion, T cells increase uptake of essential metabolic nutrients that fuel macromolecule biosynthetic processes. It is now appreciated that limiting availability of glucose^31–33^, glutamine^10,34^, alanine^13^, leucine^35^, methionine^36^ and arginine^11,37–38^ can lead to deficits in T cell activation and function. Our studies extend this understanding by revealing that extracellular Asn availability is essential for optimal activation and proliferation of helper CD4^+^ T cells. This dependency is tightly linked to protein synthesis. By targeting this metabolic dependency, we show that Asn depletion can be used to ameliorate disease severity during autoimmunity driven by pTH17 cells. This is effective whether extracellular Asn is depleted prophylactically or later during active EAE. Our observations suggest that Asn deprivation could potentially restrict the function of pathogenic effector T cells in inflammatory and autoimmune disorders. In line with our findings, others have shown that amino acid deficiency can influence Th17 cell differentiation and EAE severity. Specifically, halofuginone, a molecule that mimics amino acid restriction by inhibiting prolyl-tRNA synthetase, blocks IL-23–induced STAT3 phosphorylation and IL-17 cytokine expression in memory Th17 cells. Halofuginone-treated memory Th17 cells exhibit reduced EAE severity in vivo, mirroring our observations in mice treated with PEG-AsnASE or Asn-deficient pTh17 cells.^39–40^

Our results build upon recent reports demonstrating the importance of extracellular Asn in the activation and proliferation of CD8^+^ T cells.^14–18^ Consistent with our observations, extracellular Asn has been shown to be critical for TCR-induced activation, proliferation and metabolic reprogramming of naive CD8^+^ T cells.^14–15^ Although upregulation of ASNS enables CD8^+^ T cells to function in the absence of extracellular Asn, ASNS-expressing CD8^+^ T cells activated in Asn-deficient media exhibit significantly lower activation, proliferation and effector molecule production compared to Asn-sufficient media.^14^ Interestingly, pharmacological inhibition of ASNS activity only modestly decreases CD8^+^ T cell function, suggesting that newly synthesized Asn has a lower impact than extracellular Asn on the initiation of CD8^+^ T cell responses.^15^ Our results also demonstrate that CD4^+^ T cells require continuous extracellular Asn for optimal activation and proliferation, despite upregulating ASNS. The relative expression levels of ASNS and timing of Asn depletion also can influence the differentiation states of CD8^+^ T cells.^17–18^ While Asn depletion early during the differentiation of CD8^+^ T cells favored the maintenance of an effector phenotype^17–18^, depletion of extracellular Asn late during CD8^+^ T cell activation promoted polarization towards a central memory phenotype.^18^ We find that Asn depletion at early stages of T helper subset polarization inhibits lineage-defining cytokine production. However, further studies are needed to examine the requirement of Asn during later stages of activation and differentiation, as well as its role in supporting the longevity of CD4^+^ helper T cell responses. In addition, it remains to be determined whether the generation of memory CD4^+^ T cells or their recall responses similarly depend on extracellular Asn availability.

How does Asn depletion impair the proliferation and functional activity of T cells? In this work, we demonstrate that Asn functions as a proteinogenic amino acid, thereby promoting CD4^+^ T cell proliferation and lineage-defining cytokine production. These results are consistent with observations in mammalian cell systems, which highlight the essentiality of Asn in the setting of glutamine deprivation.^41–42^ Studies have shown that simultaneous glutamine and Asn depletion cripples T cell activation and proliferation, similar to the effects seen when ASNS-deficient T cells are deprived of extracellular Asn.^14^ Asn essentiality in glutamine deficient conditions is also reflected by their shared role as amino acid exchange factors. In a cell model of liposarcoma, Asn and glutamine export promoted import of amino acids crucial for one-carbon metabolism and mTOR activation.^41^ Since both mTOR activation and one-carbon metabolism are upregulated after TCR stimulation, it is possible that Asn-mediated amino acid exchange activity contributes an additional functional role in supporting CD4^+^ T cell activation and proliferation. Our work shows that Asn availability is important for the maintenance of mitochondrial mass and membrane potential, and its depletion promotes accumulation of depolarized mitochondria with a phenotype similar to those observed in exhausted CD8^+^ T cells.^23^ Given that Asn can directly bind to and enhance the activity of LCK^15^, a key TCR signaling kinase, it is possible that Asn might also interact with other proteins related to mitochondrial respiration. Future studies determining the interactome of Asn in activated T cells will be crucial for uncovering novel regulatory functions of Asn beyond protein synthesis.

Our finding that Asn deprivation can ameliorate the severity of EAE, even after the priming of CNS-reactive CD4^+^ T cells, warrants further investigation in models of autoimmunity and immune dysregulation. It would be interesting to explore whether therapeutic asparaginase could be utilized to prevent the induction or severity of colitis or delay the spontaneous development of diabetes in NOD mice mediated by islet-reactive CD4^+^ and CD8^+^ T cells. Future studies should investigate whether targeted delivery of asparaginases to tissues with autoreactive T cells, such as the inflamed synovium in arthritis, can similarly ameliorate autoimmunity without compromising anti-pathogen immune response essential for the host. In addition, Asn depletion may be beneficial in managing life-threatening immune-related adverse events that some cancer patients develop with immune checkpoint blockade.

In summary, our studies reveal that extracellular Asn availability represents a metabolic vulnerability for the activation and differentiation of naive CD4^+^ T cells, largely due to the requirement for Asn in sustaining protein synthesis. Therapeutic Asn depletion by targeted asparaginase treatment may provide a conceptually novel strategy for autoimmunity.

## MATERIALS AND METHODS

### Mice

Wild-type (WT) C57BL/6J and 2D2 T cell receptor (TCR) transgenic female mice were purchased from the Jackson Laboratories. 8–12-week-old female mice were used for all experiments. All mice were maintained under the guidelines and policies set by the Harvard Medical School Standing Committee on Animals and the National Institutes of Health. All mouse protocols were approved by the Harvard Medical Area Standing Committee on Animals.

### Isolation and activation of murine CD4^+^ T cells

Naive CD4^+^CD62L^+^CD44^-^ T cells were isolated from mouse spleens using the Naive CD4^+^ T Cell Isolation Kit (Miltenyi Biotec) in accordance with the manufacturers’ protocols. Following isolation, naive CD4^+^ T cells were stimulated with plate-bound anti-CD3 and anti-CD28 mAbs (Thermo Fisher Scientific) at a concentration of 4 μg/mL using a 96-well flat-bottom plate (10 x 10^4^ cells/well). For proliferation studies, naive CD4^+^ T cells were labeled with cell trace violet dye (Thermo Fisher Scientific) in accordance with the manufacturer’s protocol. Cells were subsequently cultured in standard RPMI-1640 supplemented with 10% heat inactivated FBS, 10 mM HEPES, 0.05 mM 2-mercaptoethanol and 1% penicillin-streptomycin or Asn-deficient RPMI (Thermo Fisher Scientific) supplemented with 10% heat inactivated FBS, 10 mM HEPES, 0.05 mM 2-mercaptoethanol and 1% penicillin-streptomycin. In some studies, standard RPMI media was treated with 10 IUs/L of PEGylated asparaginase (PEG-AsnASE) to remove Asn and used as an additional control. In studies in which amino acids were depleted, RPMI without amino acids (US Biological Sciences) was supplemented with 10% heat inactivated FBS, 10 mM HEPES, 0.05 mM 2-mercaptoethanol and 1% penicillin-streptomycin, with pH adjustment to 7.3, and specific amino acids were added depending on desired amino acid composition. Concentrations for each added amino acid were based on Thermo Fisher Scientific RPMI Formulation (https://www.thermofisher.com/us/en/home/technical-resources/media-formulation.114.html). All amino acids apart from threonine (Thermo Fisher Scientific) were purchased from Sigma-Aldrich. For additional activation studies, DMEM with glutamine (Thermo Fisher Scientific) was supplemented with 10% heat inactivated FBS and 1% penicillin-streptomycin and individual non-essential amino acids depending on desired condition. Prior to surface, intracellular and metabolic dye staining, cells were transferred to a 96 well V-bottom plate.

### CD4^+^ helper T cell differentiation and intracellular staining

Naive CD4^+^CD62L^+^CD44^-^ cells were isolated from mouse spleens using the Naive CD4^+^ T Cell Isolation Kit (Miltenyi Biotec) in accordance with the manufacturer’s protocols. Following isolation, naive CD4^+^ T cells were stimulated with plate-bound anti-CD3 and anti-CD28 mAbs (Thermo Fisher Scientific) at a concentration of 4 μg/mL in a 96-well flat-bottom plate (10 x 10^4^ cells/well) and cultured in either RPMI, Asn-deficient RPMI or RPMI treated with 10 IUs/L PEG-AsnASE. In some studies, 10 IUs/L PEG-AsnASE was added to cultures at 0, 6, 12, 24, 36 hours following stimulation. Naive CD4^+^ T cells were differentiated into distinct helper T (T_H_) cell subsets using the following polarization conditions and recombinant proteins and antibodies: For T_H_1 differentiation: 10 ng/mL IL-12 (Peprotech), 5 ng/mL IL-2 (Peprotech) and 10 μg/mL anti-IL-4 (Biolegend Clone 11B11). For non-pathogenic T_H_17 conditions: 20 ng/mL IL-6 (Peprotech), 2 ng/mL TGF-β1 (Peprotech), 10 μg/mL anti-IFN-γ (Biolegend Clone XMG1.2), 10 μg/mL anti-IL-4 (Biolegend Clone 11B11) and 10 μg/mL anti-IL-2 (Biolegend Clone JES6-1A12). For pathogenic T_H_17 conditions: 20 ng/mL IL-6 (Peprotech), 10 ng/mL IL-23 (R&D Systems), 10 ng/mL IL-1β (Peprotech), 10 μg/mL anti-IFN-γ (Biolegend Clone XMG1.2), 10 μg/mL anti-IL-4 (Biolegend Clone 11B11) and 10 μg/mL anti-IL-2 (Biolegend Clone JES6-1A12). For T_H_2 differentiation conditions: 40 ng/mL IL-4 and 10 μg/mL anti-IFN-γ (Biolegend Clone XMG1.2). For iTreg differentiation conditions: 2.5 ng/mL TGF-β1 (Peprotech), 10 μg/mL anti-IL-4 (Biolegend Clone 11B11), and 10 μg/mL anti-IFN-γ (Biolegend Clone XMG1.2). On day 3 cells were re-stimulated using a 1X eBioscience™ Cell Stimulation Cocktail (plus protein transport inhibitors) for 4 hours at 37°C and transferred to a 96 well V-bottom plate for intracellular staining. Cells were then washed two times using a 1X cell stain buffer solution (Biolegend) and stained with CD4 (Biolegend Clone RM4-5) and a 1:2000 fixable viability stain 780 (BD) for 30 minutes on ice, followed by two washes using 1X cell stain buffer. Intracellular staining was performed using the BD Cytofix/Cytoperm™ Fixation/Permeabilization Kit (BD) according to the manufacturer’s instructions. The following antibodies were diluted at 1:200 in 1X Perm buffer and used for intracellular staining: IL17A (Biolegend Clone TC11-18H10.1) and IFN-γ (Biolegend Clone XMG1.2). Acquisition was performed on a FACSymphony cytometer with DIVA software (BD), and data were analyzed using FCS Express Software (De Novo).

### Surface/Intracellular Staining and Flow Cytometry

Primary mouse cells were isolated from spleen and central nervous system (CNS) including brain and spinal cord. Single-cell suspensions were incubated with 1:100 TruStain FcX™ (Biolegend) 1X DPBS solution for 15 minutes at room temperature to block Fc receptors. Viability was assessed using a fixable viability stain 780 (BD Biosciences) at a 1:1000 dilution in 1X DPBS for 20 minutes on ice followed by one wash in cell stain buffer (Biolegend). For surface staining, cell suspensions were incubated using cell stain buffer (Biolegend) and brilliant stain buffer (BD) at a 1:1 ratio for 30 minutes on ice in the dark followed by 2 washes with cell stain buffer (Biolegend). Cells were then resuspended in a 1X stabilizing fixative (BD) solution. Intracellular staining was performed using the BD Cytofix/Cytoperm™ Fixation/Permeabilization Kit (BD) according to the manufacturer’s instructions. The following antibodies were used: anti-CD4 (Biolegend, RM4-5), CD25 (Biolegend, PC61), CD3 (BD Biosciences, 145-2C11), CD69 (Biolegend, H1.2F3), CD44 (Biolegend, IM7), CD71 (Biolegend, RI7217), PD-1 (Biolegend, 29F.1A12), anti-Foxp3 (Ebioscience, FJK-16s), anti-IL17A (Biolegend, TC11-18H10.1), IFNγ (Biolegend, XMG1.2), TCRβ (BD Biosciences, H57-597), GATA3 (Biolegend, 16E10A2), RORyT (eBioscience, B2D), CD8 (BD Biosciences, 53-6.7), CD45 (BD Biosciences, 30F11), Ki-67 (BD Biosciences, B56), GM-CSF (Biolegend, MP1-22E9), IL-22 (Biolegend, Poly5164), IL17F (Biolegend, 9D3.1C8), Tbet (Biolegend, 4B10). Annexin-V staining was done using the FITC Annexin V Apoptosis Detection Kit I (BD) in accordance with the manufacturer’s instructions. Acquisition was performed on a FACSymphony cytometer with DIVA software (BD), and data were analyzed using FCS Express Software (De Novo).

### Mitochondrial and metabolic dye staining

Naive CD4^+^CD62L^+^CD44^-^ cells were isolated from mouse spleens using the Naive CD4^+^ T Cell Isolation Kit (Miltenyi Biotec) according to the manufacturer’s instructions. Following isolation naive CD4^+^ T cells were stimulated with plate-bound anti-CD3 and anti-CD28 mAbs (Thermo Fisher Scientific) at a concentration of 4 μg/mL in a 96-well flat-bottom plate (10 x 10^4^ cells/well) and cultured in either RPMI, Asn-deficient RPMI or RPMI treated with 10 IUs/L PEG-AsnASE. For Mitotracker^TM^ green (Thermo Fisher Scientific) and tetramethyl rhodamine, Methyl Ester, Perchlorate (Thermo Fisher Scientific) staining, cells were subsequently incubated at 37°C for 30 minutes in 200 μL of prewarmed RPMI media containing 100 nM MTG and TMRM followed by two washes in 1X DPBS. Viability and cell surface staining was performed as described above.

### Seahorse analysis

The Oxygen Consumption Rate (OCR) and Extracellular Acidification Rate (ECAR) were evaluated under mitochondrial and glycolysis stress test conditions, respectively, following the manufacturer’s instructions and protocols, utilizing a XFe96 Extracellular Flux Analyzer (Agilent). One day prior to the measurement, an Agilent Seahorse XFe96 Sensor Cartridge (Agilent) was hydrated with HPLC grade water in a CO2-free incubator. On the subsequent day, the solution was replaced with XF Calibrant (Agilent), and the cartridge was maintained in a 37°C CO2-free incubator for a minimum of two hours. Naive CD4^+^ T cells were stimulated with anti-CD3 and anti-CD28 mAbs (Thermo Fisher Scientific) at a concentration of 4 μg/mL in a flat-bottom 48-well plate for 2 days, using either RPMI, Asn-deficient RPMI, or RPMI treated with 10 IUs/mL PEG-AsnASE at 0, 6, 12, 24, 36 hours following stimulation. After 48 hours, CD4^+^ T cells were enumerated and transferred to a poly-D-lysine-coated Seahorse XF96 tissue culture microplate (Agilent) at a density of 100,000 cells/well. Seahorse XP RPMI or DMEM medium (Agilent), comprising, 2 mM L-glutamine, and 1 mM sodium pyruvate, was used for assessment of ECAR and 10 mM glucose was added. In some experiments, asparaginase (AsnASE) from E. coli (Sigma-Aldrich) was injected first for a final volume of 10 mM under mitochondrial stress test conditions. In additional experiments, naive CD4^+^ T cells were stimulated with anti-CD3 and anti-CD28 mAbs at a concentration of 4 μg/mL in a flat-bottom 6-well plate for 1 day supplemented with either asparagine, alanine, glutamate, aspartate, or proline at a 0.38 mM final concentration. After 24 hours, CD4^+^ T cells were processed as described above.

### Chemicals

^15^N_2_-L-Asn hydrate [Chemical Formula H2*NCOCH2CH*(NH2)COOH:H20] was acquired from Cambridge Isotope Laboratories Inc (Cat # NLM-3286-0) with a documented purity ≥ 98% as determined HPLC. For tracing studies, ^15^N_2_-L-Asn hydrate was dissolved in Asn-deficient RPMI media at a final concentration of 0.38 mM. All PEG-AsnASE experiments were performed using pegaspargase (Oncaspar, Shire Pharmaceuticals, Lexington, MA), an FDA-approved PEGylated form of E. coli asparaginase.

### Immunoblotting

Equal numbers of cells were washed once with 1X DPBS and lysed by adding 1X SDS sample buffer (Sigma) with subsequent boiling for 15 minutes. The resulting cell extracts were clarified via centrifugation at 13,000 x g, separated through SDS-PAGE, and then transferred to nitrocellulose membranes (Bio-Rad) using electrophoresis. Membranes were then immersed in Tris-buffered saline (TBST) buffer with 3% (w/v) bovine serum albumin (BSA) for a 30-minute blocking period followed by incubation with ASNS primary antibody (Cell Signaling Technology) diluted in blocking buffer overnight at 4°C. Membranes were washed three times for 1 hour and subsequently incubated with anti-mouse IgG HRP-conjugated secondary antibody (Thermo Fisher Scientific) for 1 hour at RT, followed by 3 washes and visualization using the Western Lightning ECL Pro <u>Chemiluminescence</u> Substrate (PerkinElmer).

### Bulk RNA sequencing analysis

The original data was obtained from <u>GSE206304</u>^21^ and reanalyzed to examine expression of Asn metabolism-related genes including *Asns*, *Asrgl1, Slc1a5, Slc38a2, and Slc6a14* in helper T cells differentiated in vitro under T helper 1 (T_H_1), non-pathogenic T helper 17 (npT_H_17), and pathogenic T helper 17 (pT_H_17) polarizing conditions. Reads were aligned to the mm10 genome using Tophat followed by duplicate removal and htseq counts were used for the generation of gene count tables. Read counts were normalized using the DESeq2 package.

### Real time PCR

Total DNA from CD4^+^ T cells was extracted using the DNA Micro kit (Qiagen). DNA was quantified using the Qubit™ dsDNA Quantification Assay Kits (Thermo Fisher Scientific). cDNA was subsequently synthesized using the iSCRIPT kit (BioRad). Quantitative PCR analysis was conducted using the SYBR Green Fast Mix (Quanta BioSciences) on a LightCycler 96 Instrument (Roche). Primers used: Asns: F: 5′- GATCTTCATCGCACTCAGACA-3′, R: 5′-CCTCTGCTCCAC CTTCTCT-3′; Asrgl1: F: 5′- GATACTTTCCCCATGTCCTGTG-3′, R: 5′-TTGGCTTACGCAACC TCTAC-3′. DNA concentrations were within the linear range of the primers.

### OPP Protein Synthesis Assay

Naive CD4^+^CD62L^+^CD44^-^ cells were isolated from mouse spleens using the Naive CD4^+^ T Cell Isolation Kit (Miltenyi Biotec) according to the manufacturer’s instructions. Following isolation, naive CD4^+^ T cells were stimulated with plate-bound anti-CD3 and anti-CD28 mAbs (Thermo Fisher Scientific) at a concentration of 4 μg/mL in a 96-well flat-bottom plate (20 x 10^4^ cells/well) and cultured in either RPMI or Asn-deficient RPMI. After 24 hours, 0.38 mM Asn was added to samples cultured in Asn-deficient RPMI and cells were cultured for an additional 4 hours. For detection of nascent protein synthesis, the Click-iT™ Plus OPP Alexa Fluor™ 488 Protein Synthesis Assay Kit (Thermo Fisher Scientific) was used in accordance with the manufacturer’s protocols. As a positive control, some cell suspensions were treated with 50 μg/mL cycloheximide (Sigma Aldrich) for 30 minutes at 37°C to block protein synthesis. Cell suspensions were then resuspended in 200 μL of a 1X stabilizing fixative (BD) solution followed by flow cytometry assessment for OPP fluorescent intensity using a FACSymphony cytometer with DIVA software (BD).

### Isolation of protein and hydrolysis into amino acid monomers

Cell pellets were resuspended in 200 μL of lysis buffer containing 2% SDS, 150 mM NaCl, 50 mM Tris (pH 8.5), proteinase inhibitor mix (Roche), 5 mM DTT, and incubated on ice for 10 minutes followed by incubation at 60°C for 45 minutes. After cooling to room temperature, iodoacetamide was added to each sample for a final concentration of 14 mM, and the samples were incubated for an additional 45 minutes. The treated samples were then mixed with a solution consisting of 3 parts ice-cold methanol, 1 part chloroform, and 2.5 parts H_2_O followed by centrifugation at 4000 x g for 10 minutes. The top layer was then removed and 3 parts of ice-cold methanol were added, followed by centrifugation at 4000 x g for 5 minutes. Following removal of the top layer, the samples were mixed with 3 parts of ice-cold acetone, vortexed, and centrifuged at 4000 x g for 5 minutes. The pellet was then washed with 2 mL of ice-cold acetone and stored at −80°C prior to chemical hydrolysis. The protein pellet obtained was resuspended in 6 N HCl/acetic acid (50:50,100 μL) and subjected to heating at 95°C for 1 hour. The resulting aqueous solution was diluted into a mixture of 40% acetonitrile, 40% methanol, and 20% water and analyzed by LC-MS.

### Induction of EAE

EAE was induced by immunization of 10-week-old female C57BL/6J mice with the Hooke Kit^TM^ MOG_35-55_/CFA Emulsion PTX in accordance with the manufacturer’s instructions. Briefly, mice were acclimated in our animal facility for at least 7 days prior to immunization. For EAE induction, mice were immunized with 200 μg of antigen (MOG_35-55_) in emulsion with complete Freund’s adjuvant (CFA) in both flanks followed by administration of 120 ng pertussis toxin (PTX) on the day of immunization and following day. Mice were monitored for signs of clinical disease and scored following the Hooke scoring system (https://hookelabs.com/services/cro/eae/MouseEAEscoring.html). For analysis of cellular infiltrates in the CNS, brain and spinal cords were isolated at the peak of disease. Prior to CNS collection, mice were perfused with 1X DPBS and brains and spinal cords were mechanically dissociated through a 70 μm nylon cell strainer followed by digestion with Collagenase D (Sigma-Aldrich) for 20 minutes in a 37°C shaker. Digests were then filtered through a 70 μm strainer, resuspended in a 30% Percoll/DPBS solution and overlaid over a 70% Percoll gradient for mononuclear cell isolation. Following centrifugation at 800 x g for 30 minutes at room temperature, lymphocytes in the interface were collected, washed with RPMI media, and stimulated using a 1X eBioscience™ Cell Stimulation Cocktail plus protein transport inhibitors (Thermo Fisher Scientific) for 4 hours at 37°C. Cells were then centrifuged at 600 x g for 3 minutes followed by resuspension in cell stain buffer (Biolegend) for subsequent flow cytometry staining.

### Pathogenic T_H_17 differentiation for EAE induction and adoptive transfer

Naive CD4^+^CD62^+^ CD44^-^ cells were isolated from the spleens of female 2D2 TCR transgenic mice using the Naive CD4^+^ T Cell Isolation Kit (Miltenyi Biotec) in accordance with the manufacturer’s protocols. 2×10^6^ naive CD4^+^ T cells were then stimulated with plate-bound anti-CD3 and anti-CD28 mAbs (Thermo Fisher Scientific) at a concentration of 4 μg/mL in a 48 well flat-bottom plate and cultured for 3 days in either RPMI media or Asn-deficient RPMI media under pathogenic T_H_17 polarizing conditions: 20 ng/mL IL-6 (Peprotech), 10 ng/mL IL-23 (R&D Systems), 10 ng/mL IL-1β (Peprotech), 10 μg/mL anti-IFN-γ (Biolegend Clone XMG1.2) and 10 μg/mL anti-IL-4 (Biolegend Clone 11B11). Cell suspensions were then rested for 2 days in the absence of TCR stimulation using either RPMI media or Asn-deficient RPMI media containing 20 ng/mL IL-23 (R&D Systems). After two days of rest, cells were restimulated with plate-bound anti-CD3 and anti-CD28 mAbs (Thermo Fisher Scientific) at a concentration of 4 μg/mL in a flat bottom 6 well plate for two days, followed by two washes in 1X DPBS. Following counting a small aliquot of cells to determine viability by flow cytometry, 4×10^6^ viable T cells from each respective culture condition were transferred by intravenous injection into C57BL/6J female recipient mice to induce EAE. Mice were monitored for signs of clinical disease and scored following the Hooke scoring system.

### Statistical Analysis

Statistics were computed with GraphPad Prism 9 software (GraphPad Software) using unpaired Student’s T-test for comparisons between two groups, one-way ANOVA followed by Tukey’s or Dunnets multiple comparison when comparing three or more groups, or Two-way ANOVA for multiple comparisons within groups. Graphs containing EAE clinical scores represent mean values with error bars representing the Standard Error of the Mean (SEM). Unless noted otherwise, all other data are represented as mean +/- SD. P-values are denoted in figures as: *p<0.05, **p<0.01, ***p<0.001, ****<0.0001.

## ACKNOWLEDGEMENTS

This work was supported by NIH P01 AI056299, P01 AI039671 and AI108545 (to A.H.S.), NIH U54 CA224088, and R01CA276866 (to A.H.S. and M.C.H.), and the Ludwig Center at Harvard Medical School, NIH U01 CA267827, and the Paul F. Glenn Foundation for Medical Research to M.C.H. P.G. was supported by a predoctoral NIH fellowship 1F31CA281090-01. S.H. was supported by the Banting postdoctoral fellowship from the Canadian Institutes of Health Research (CIHR). K.K. is a Gilead Sciences Fellow of the Life Sciences Research Foundation. Schematics were created using BioRender.com. We would like to thank members of the Sharpe and Haigis laboratories for productive discussion.

## AUTHORS CONTRIBUTIONS

**P.G**: Conceptualization, resources, data curation, formal analysis, validation, investigation, visualization, methodology, writing-original draft, writing review and editing, funding acquisition. **S.J**: data curation, formal analysis, validation, investigation, visualization, methodology, writing-original draft, writing review and editing. **K.K**: Conceptualization, data curation, formal analysis, writing review and editing. **S.H**: data curation, formal analysis. **D.P:** methodology, data curation, resources. **D.L:** methodology, data curation. **S.K:** methodology, data curation. **J.R** advising, supervision, resources, methodology. **A.H.S** and **M.C**: Conceptualization, resources, supervision, funding acquisition, project administration, writing-review and editing.

---

**Supplemental Figure 1.**
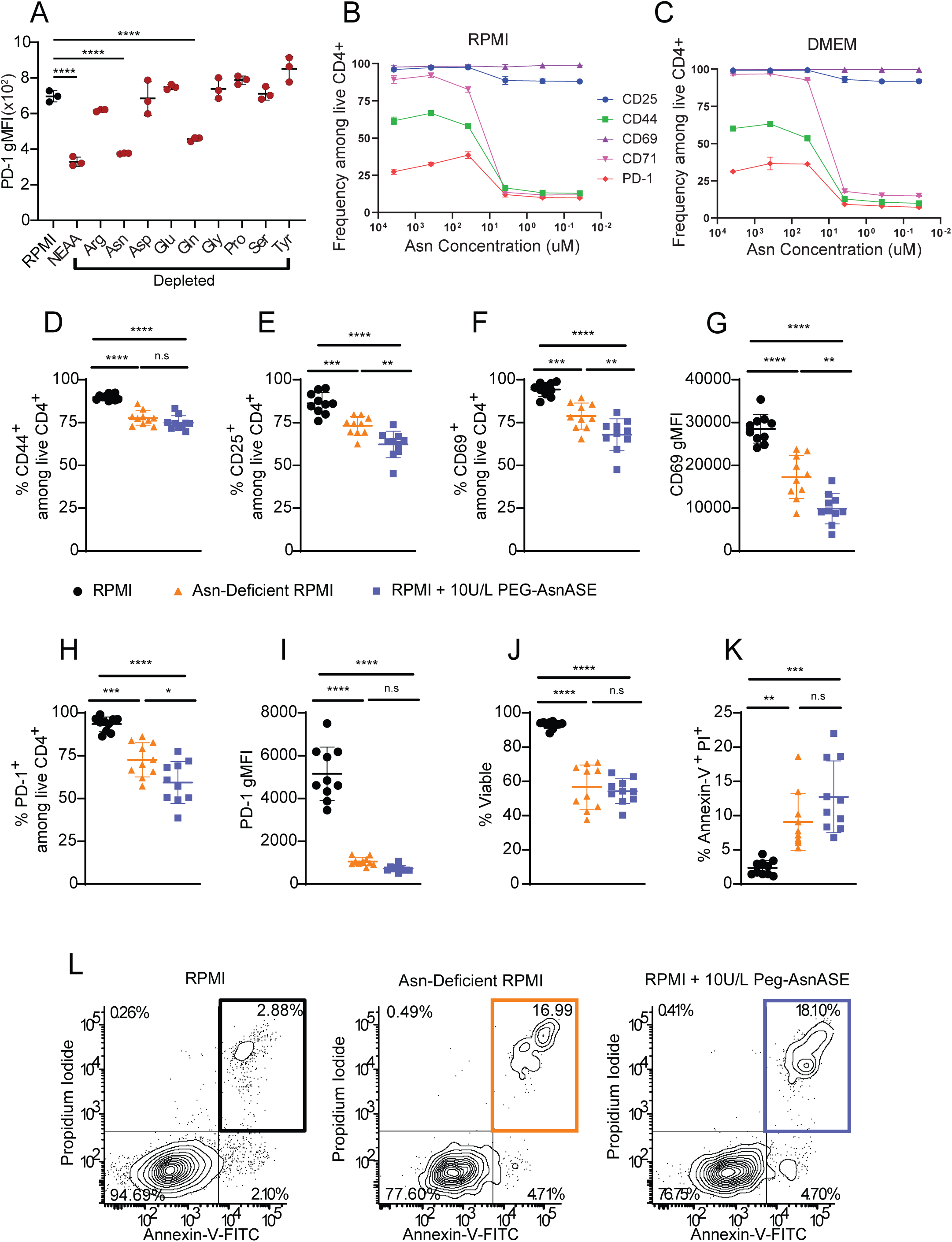
Extracellular asparagine is essential for CD4^+^ T cell activation. (**A**) Expression levels of the activation marker PD-1 following 24 hours of stimulation with plate-bound anti-CD3/CD28 mAbs in RPMI lacking the indicated individual amino acids shown in red. Non-essential amino acids (NEAA) include asparagine (Asn), aspartate (Asp), glutamate (Glu), proline (Pro), arginine (Arg), glutamine (Gln), glycine (Gly), serine (Ser), and tyrosine (Tyr). (**B-C**) Frequency of cells expressing the activation markers CD25, CD44, CD69, CD71, and PD-1 following 24 hours of stimulation with plate-bound anti-CD3/CD28 mAbs in RPMI without Asn (**B**) or DMEM (**C**) supplemented with increasing concentrations of Asn. Quantification of the proportions of CD4^+^ T cells expressing cell surface activation markers CD44 (**D**), CD25 (**E**), CD69 (**F**), and PD-1 (**H**) as well as expression levels of CD69 (**G**) and PD-1 (**I**) on a per cell basis following 24 hours of stimulation with plate-bound anti-CD3/CD28 mAbs in complete RPMI (RPMI), Asn-deficient RPMI or RPMI with 10 IUs/L PEGylated-asparaginase (PEG-AsnASE) added at the start of culture. (**J**) Quantification of the proportions of viable CD4^+^ T cells in each respective culture condition after 3 days of stimulation with plate-bound anti-CD3/CD28 mAbs in RPMI, Asn-deficient RPMI or RPMI with 10 IUs/L PEG-AsnASE added at the start of culture. (**K**) Quantification of the proportion of Annexin-V propidium iodide double-positive CD4^+^ T cells as shown in L. (**L**) Representative flow cytometry contour plots depicting Annexin-V and propidium iodide staining in naive CD4^+^ T cell following 2 days of stimulation with plate-bound anti-CD3/CD28 mAbs in RPMI, Asn-deficient RPMI or RPMI with 10 IUs/L PEG-AsnASE added from the initiation of culture. Upper right quadrants depict the proportions of CD4^+^ T cells that have undergone apoptosis in each respective culture condition. Each dot represents cells obtained from an individual animal (D-K). Results are shown as mean ± SD and are pooled from 2 independent experiments (D-K) or a representative of 2 independent experiments (A-C, L). non-significant (n.s.), *p < 0.05 **p < 0.01, ***p < 0.001, ****p < 0.0001, one-way ANOVA with Turkey’s multiple comparison test (D-K) or Dunnet’s multiple comparison test (A).

**Supplemental Figure 2.**
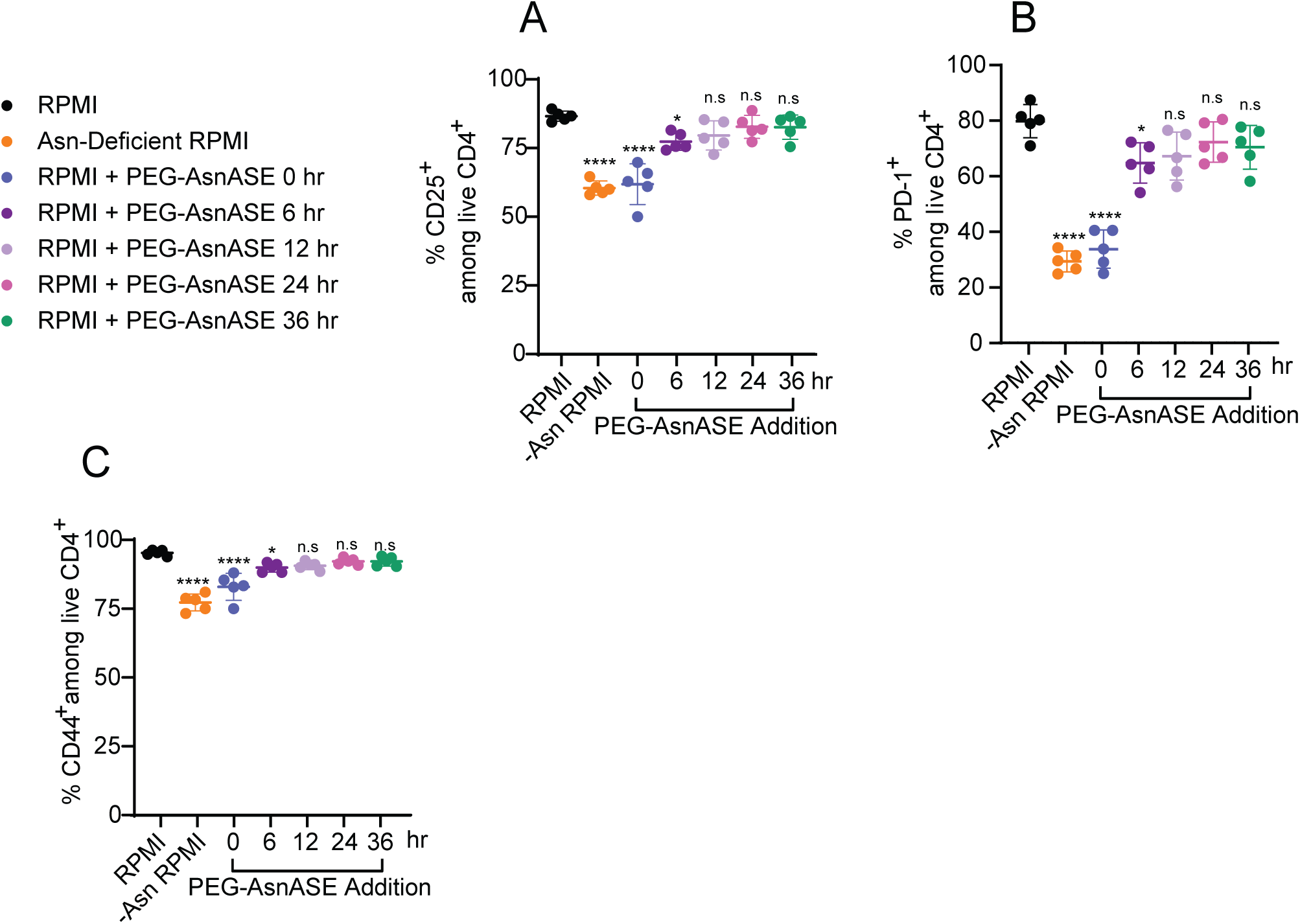
Expression of CD4^+^ T cell activation proteins in asparagine sufficient and deficient conditions. (**A-C**) Quantification of the proportions of CD4^+^ T cells expressing CD25 (**A**), PD-1 (**B**), and CD25 (**C**) following 2 days of stimulation with plate-bound anti-CD3/CD28 mAbs in complete RPMI, Asn-deficient RPMI or RPMI with 10 IUs/L PEGylated-asparaginase added at 0, 6, 12, 24, or 36 hours. Each dot represents cells obtained from an individual animal (A-C). Results are shown as mean ± SD and are representative of 2 independent experiments. non-significant (n.s.), *p < 0.05, ****p < 0.0001, one-way ANOVA with Dunnet’s multiple comparison test.

**Supplemental Figure 3.**
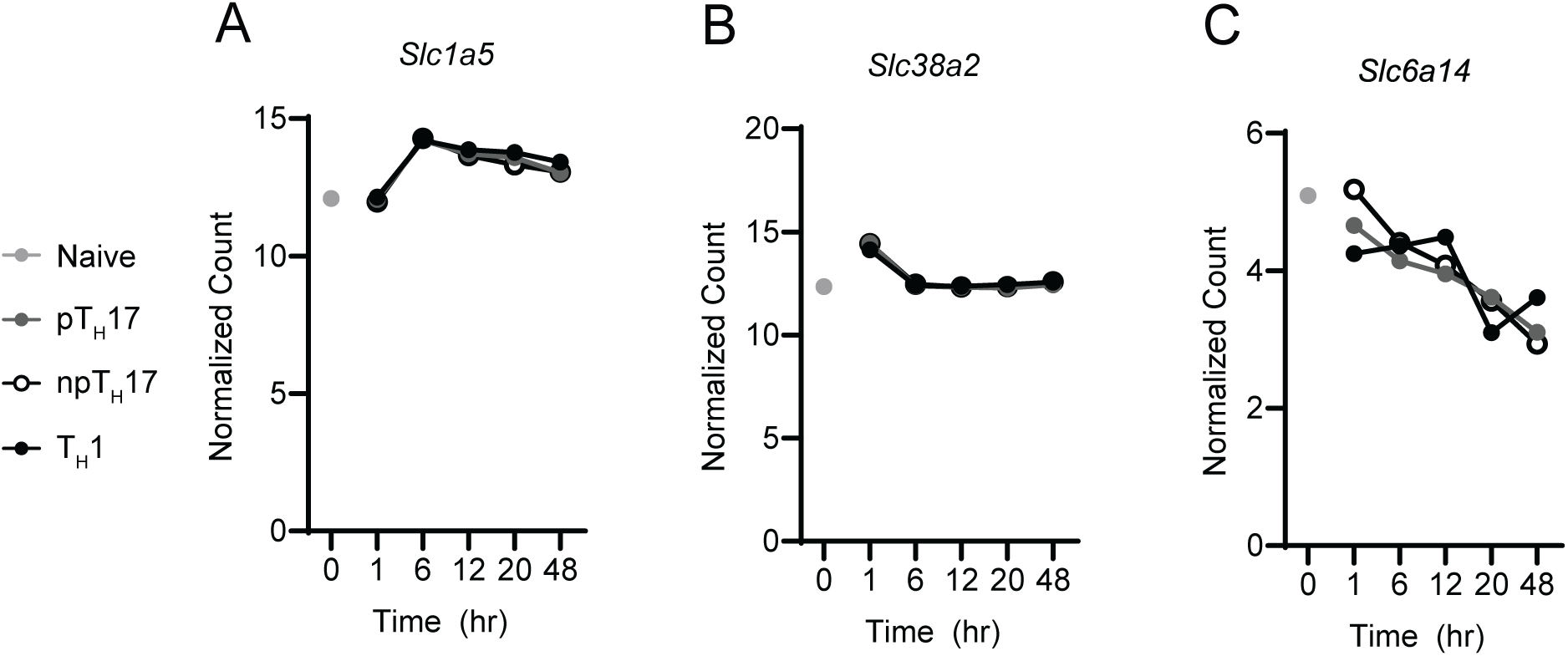
Asparagine amino acid transporters are expressed in activated CD4^+^ T cells. (**A-C**) Bulk RNA-seq analysis showing expression of *Slc1a5* (**A**), *Slc38a2* (**B**), and Slc6a14 (**C**) in naive CD4^+^CD62L^+^CD44^-^ T cells at baseline (gray dot) or following stimulation with anti-CD3/CD28 mAbs under T helper 1 (T_H_1), pathogenic T helper 17 (pT_H_17) and non-pathogenic T helper 17 (npT_H_17) conditions for 1, 6, 12, 20 and 48 hours. Results are shown as average (n=3 for each condition, per time point).

**Supplemental Figure 4.**
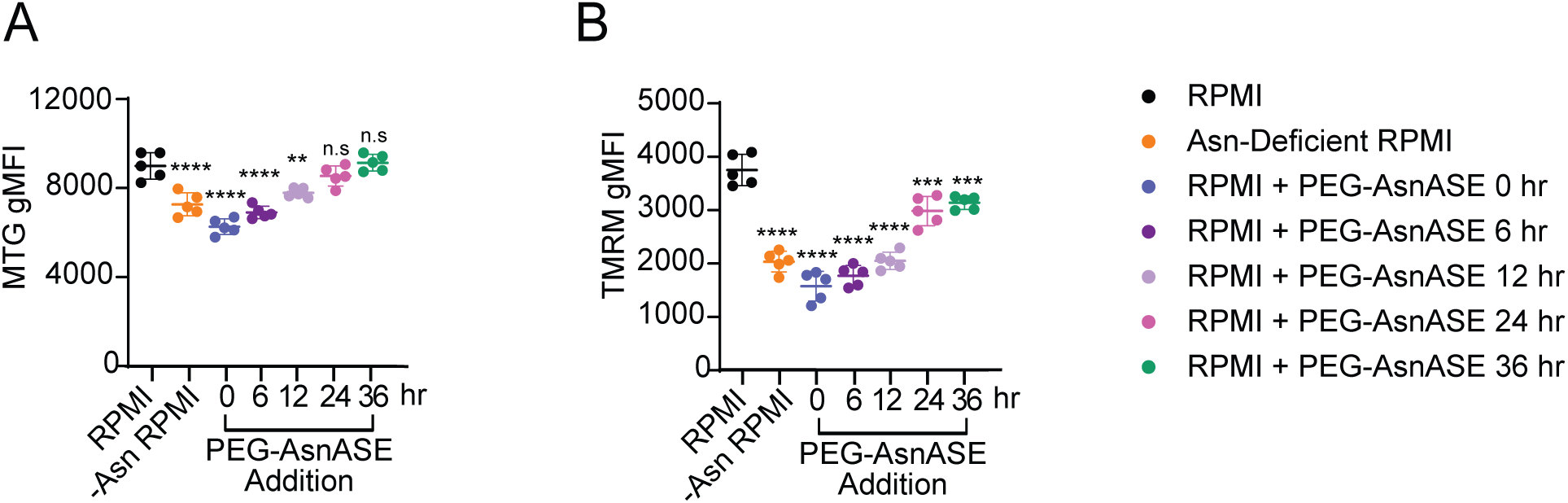
Asparagine deprivation impairs mitochondrial function in CD4^+^ T cells. (**A-B**) Naive CD4^+^ T cells were stimulated for 48 hours with plate-bound anti-CD3/CD28 mAbs in either complete RPMI media (RPMI), Asn-deficient RPMI or RPMI treated with 10 IUs/L PEGylated-asparaginase (PEG-AsnASE) added at 0, 6, 12, 24, or 36 hours. (**A**) Quantification of mitotracker green (MTG) gMFI. (**B**) Quantification of tetramethyl rhodamine methyl ester (TMRM) gMFI. Results are shown as mean ± SD and are representative of at least 2 independent experiments. non-significant (n.s.), **p < 0.01, ***p < 0.001, ****p < 0.0001, one-way ANOVA with Dunnet’s multiple comparison test.

**Supplemental Figure 5.**
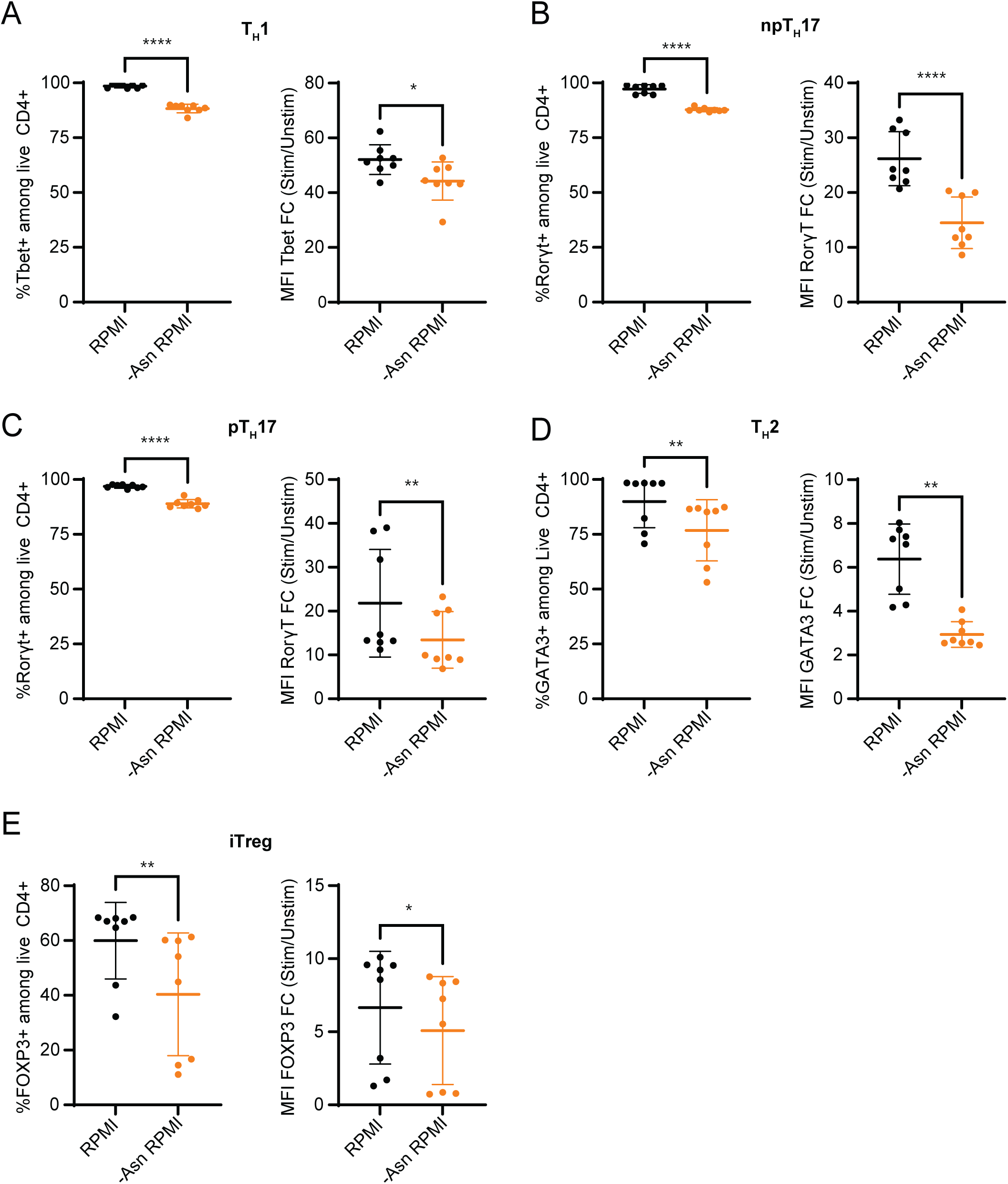
Asn depletion impairs CD4 T cell transcription factor expression. (**A-E**) Purified naive CD4^+^ T cells were activated in vitro with plate-bound anti-CD3/CD28 mAbs under T helper 1 (T_H_1), pathogenic T helper 17 (pT_H_17), non-pathogenic T helper 17 (npT_H_17), T helper 2 (T_H_1), and induced T regulatory cell (iTreg) conditions in either complete RPMI (RPMI) media or Asn-deficient RPMI. On day 3 cells were restimulated for 4 hours with phorbol 12-myristate 13-acetate (PMA), ionomycin, brefeldin A and monensin for intracellular staining. (**A**) Quantification of the proportions of Tbet-expressing CD4^+^ T cells under T_H_1 differentiation conditions and TBET gMFI. (**B**) Quantification of the proportions of RORγT-expressing CD4^+^ T cells under npT_H_17 differentiation conditions and RORγT gMFI. (**C**) Quantification of the proportions of RORγT-expressing CD4^+^ T cells under pT_H_17 differentiation conditions and RORγT gMFI. (**D**) Quantification of the proportions of GATA3-expressing CD4^+^ T cells under T_H_2 differentiation conditions and GATA3 gMFI. (**E**) Quantification of the proportions of FOXP3-expressing CD4^+^ T cells under iTreg differentiation conditions and FOXP3 gMFI. Each dot represents cells obtained from an individual animal. Results are shown as mean ± SD and are pooled from 2 independent experiments. *p < 0.05 **p < 0.01, ****p < 0.0001, Student’s T Test.

**Supplemental Figure 6.**
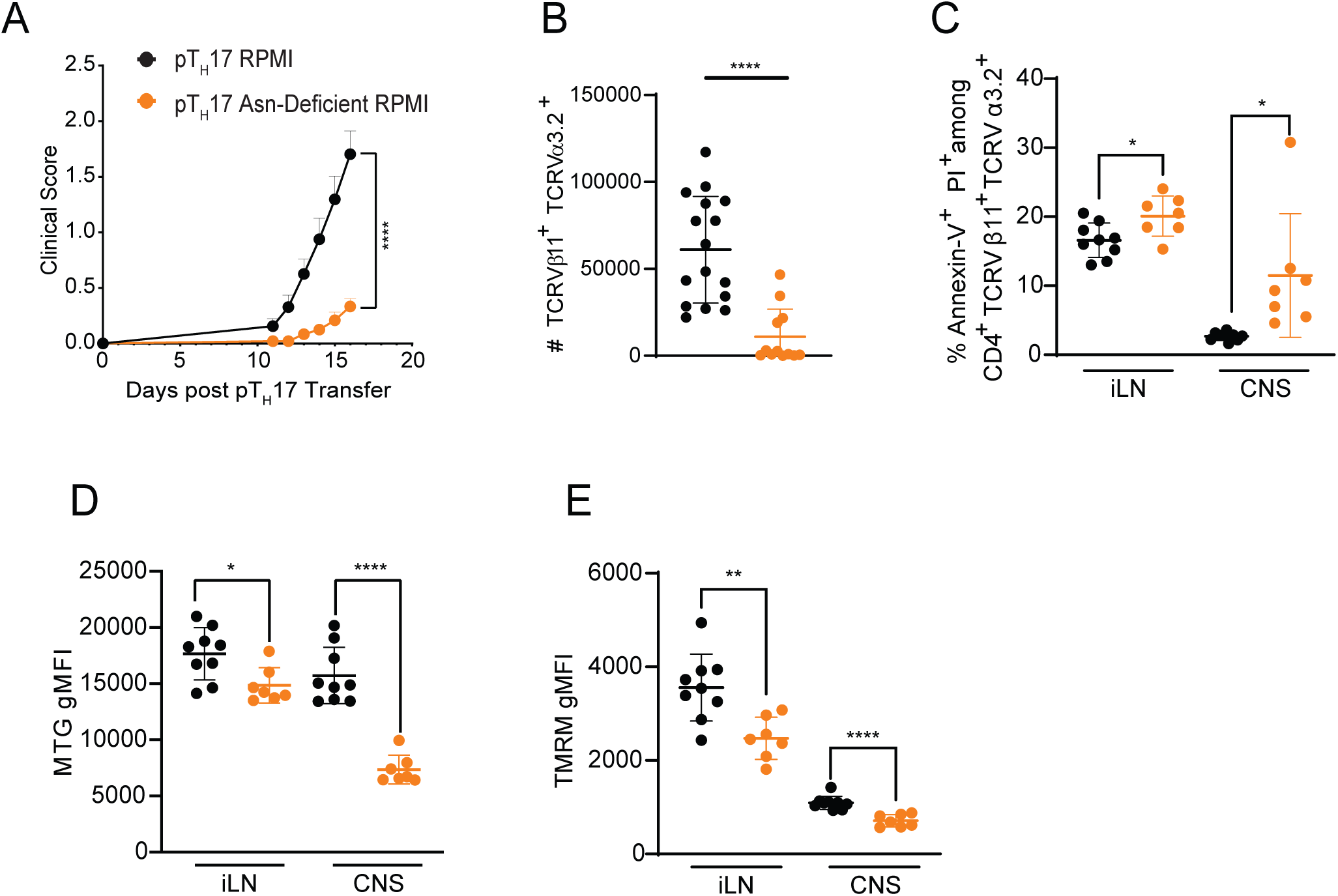
Asparagine deprivation impairs pT_H_17 function in a model of induced EAE. (**A)** Pathogenic T helper 17 (pT_H_17) cells were differentiated from naive CD4^+^ FoxP3^-^ T cells from 2D2 TCR transgenic mice in RPMI media with or without Asn. Viable pT_H_17 polarized 2D2 cells were adoptively transferred (4×10^6^/mouse) into 10-week-old C57BL/6J female recipients to induce EAE. Mice were scored daily for signs of disease (pT_H_17 RPMI n=16, pT_H_17 Asn-deficient RPMI n=12). (**B**) At peak EAE (Day 16), 2D2 cells were isolated from the CNS and inguinal lymph (iLN) node and analyzed. Quantification of the absolute numbers of central nervous system (CNS)-infiltrating Vβ11^+^Vα3.2^+^ 2D2 pT_H_17 cells at peak of EAE. (**C**) Quantification of the proportions of Vβ11^+^Vα3.2^+^ 2D2 pT_H_17 cells that have undergone apoptosis (Annexin-V^+^PI^+^) in the CNS and iLN node at peak of EAE. (**D-E**) Mitotracker green (MTG) and tetramethyl rhodamine methyl ester (TMRM) gMFI in iLN and CNS infiltrating TCRVβ11^+^TCRVα3.2^+^ CD4^+^ T cells at peak of EAE. Each dot represents an individual mouse (B-E). Results are shown as mean ± SD and are pooled (A-B) or a representative of at least 2 independent experiments (C-E). non-significant (n.s.), *p < 0.05 **p < 0.01, ****p < 0.0001, two-way ANOVA (A), and Student’s t-test (B-E).

